# Motor Learning Promotes Remyelination via New and Surviving Oligodendrocytes

**DOI:** 10.1101/2020.01.28.923656

**Authors:** Clara M. Bacmeister, Helena J. Barr, Crystal R. McClain, Michael A. Thornton, Dailey Nettles, Cristin G. Welle, Ethan G. Hughes

**Affiliations:** Department of Cell and Developmental Biology, University of Colorado School of Medicine; Department of Neurosurgery, University of Colorado School of Medicine; Department of Physiology and Biophysics, University of Colorado School of Medicine

## Abstract

Oligodendrocyte loss in neurological disease leaves axons vulnerable to damage and degeneration, and activity-dependent myelination may represent an endogenous mechanism to improve remyelination following injury. Here, we report that while learning a forelimb reach task transiently suppresses oligodendrogenesis, it subsequently increases OPC differentiation, oligodendrocyte generation, and retraction of pre-existing myelin sheaths in the forelimb region of motor cortex. Immediately following demyelination, motor cortex neurons exhibit hyperexcitability, motor learning is impaired, and behavioral intervention provides no long-term benefit to remyelination. However, partial remyelination restores neuronal and behavioral function. Motor learning following partial remyelination increases oligodendrogenesis and enhances the ability of mature oligodendrocytes to generate new myelin sheaths, resulting in almost double the remyelination of denuded axons relative to untrained controls. Together, our findings demonstrate that the correct timing of behaviorally-induced neuronal circuit activation improves recovery from demyelinating injury via enhanced remyelination from new and surviving oligodendrocytes.

---

Oligodendrocytes, the myelin-forming cells of the central nervous system (CNS), enhance the propagation of axonal action potentials and support neuronal and axonal integrity through metabolic coupling^1,2^. Injury to oligodendrocytes critically affects axonal health and is associated with significant neurologic disability in patients with multiple sclerosis (MS)^3^. While a growing number of immunotherapies decrease the frequency of MS attacks, they do not fully prevent axonal degeneration nor the accumulation of disability^4^. Oligodendrocyte precursor cells (OPCs) can generate new oligodendrocytes with the capacity to remyelinate denuded axons, which can restore neuronal function^5,6^. However, remyelination is typically incomplete in patients with MS^7^ and approaches to increase myelin repair remain limited^8,9^. Interestingly, immense variability has been observed in rates of remyelination of individual MS patients^10^, suggesting the existence of endogenous mechanisms that may modulate myelin repair following demyelinating injury.

Motor learning has been shown to drive white matter changes in both humans and rodents, in part by eliciting the proliferation and differentiation of OPCs in the adult CNS^11–13^. Similar to the context of learning, OPCs respond to demyelinating injuries with increased proliferation and differentiation^5,14^, yet it remains unclear whether learning during demyelination has synergistic or antagonistic effects. Behavioral interventions are increasingly implementable, personalizable, and measurable in clinical settings with the advent of new technologies, and have already been used to ameliorate motor function in myelin disease patients^15^. Optimizing the modality and timing of behavioral interventions may allow endogenous oligodendrogenesis-stimulating mechanisms to act in synchrony and drive more robust remyelination responses following injury in a patient-specific manner.

Regenerative processes, such as remyelination, are generally thought to respond similarly to a wide array of injuries^16^. The cuprizone-mediated demyelination model results in cortical demyelination with similar features to cortical lesions found in MS patients, notably ongoing oligodendrocyte death and regeneration^17^. We utilized this model to investigate the fundamental biology underlying remyelination without the confounding influences of autoimmunity. Through longitudinal *in vivo* two-photon imaging of oligodendrocyte lineage cells and individual myelin sheaths, we defined the complex dynamics between motor skill acquisition and its effects on oligodendroglia in both developmental and remyelinating contexts in the intact motor cortex. We found that motor learning, when properly timed, could enhance oligodendrogenesis after injury and recruit mature oligodendrocytes to participate in remyelination.

## Results

### Forelimb reach training dynamically modulates oligodendrocyte lineage cells and myelination

Previous assessments of adult oligodendrogenesis using cell lineage tracing^13,18^ and stage specific markers^19^ indicate that neuronal activity and motor skill learning rapidly increase the generation of new oligodendrocytes and myelin sheaths. However, the cellular dynamics of activity-dependent myelination remain unclear due to both incomplete labeling of the population of differentiating OPCs and inter-individual variability inherent to cross-sectional approaches. To determine the dynamics of oligodendrocytes, myelin, and OPCs during learning, we used longitudinal two-photon *in vivo* imaging in the forelimb-region of motor cortex throughout learning and rehearsal of a skilled, single-pellet contralateral forelimb reach task^20^ (**Fig. 1a,b**; **Supplementary Fig. 1; Supplementary Video 1**). To examine the effects of learning on the rate of oligodendrogenesis and remodeling of pre-existing myelin sheaths in healthy mice, we used transgenic mice that express EGFP in mature oligodendrocytes (*MOBP-EGFP*; **Fig. 1c**), which allow the visualization of all myelinating oligodendrocytes in the cortex as well as individual myelin sheaths^21^ (**Supplementary Video 2**). Long-term *in vivo* imaging of layers 1-3 of the forelimb region of motor cortex allowed us to track ~100 myelinating oligodendrocytes and their associated myelin sheaths over the course of 2-3 months per mouse. We confirmed that EGFP in *MOBP-EGFP* transgenic mice faithfully reflects the presence and length of myelin sheaths by immunohistochemistry, *in vivo* SCoRe imaging, and semi-automated tracing using a hessian-based filter algorithm (Simple Neurite Tracer^22^; **Supplementary Fig. 2**). Furthermore, using semi-automated tracing of *in vivo* z-stacks from *MOBP-EGFP* mice, we traced myelin sheaths and their connecting processes to the originating oligodendrocyte cell body (**Supplementary Fig. 3; Supplementary Video 3**). Assuming a normal range of 45±4 sheaths per cortical oligodendrocyte^21^, our average sampling was 67-74% of all sheaths on an individual oligodendrocyte.

**Fig. 1.**
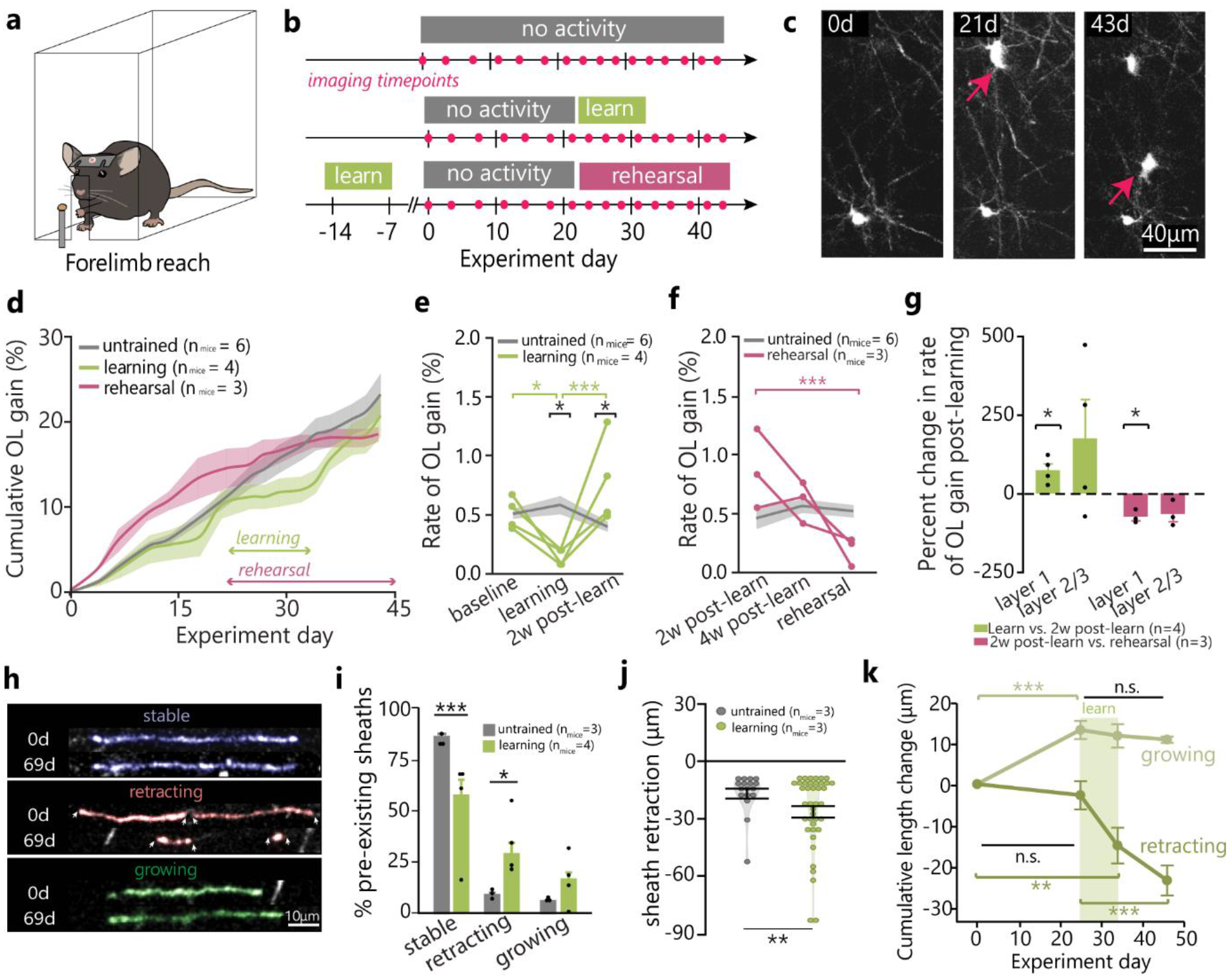
Forelimb reach training modulates oligodendrogenesis and remodeling of pre-existing myelin sheaths. **a, b,** Illustration and imaging timeline of behavioral interventions. **c,** Example of motor cortex oligodendrogenesis; red arrows indicate new cells. **d,** Cumulative oligodendrogenesis (% increase from baseline; Mean±SEM) by group. **e,** Learning modulates oligodendrogenesis rate (F_2,16_=15.61, p=0.0002; grey line±shaded area represents control Mean±SEM; green traces represent individual learning mice). Rate is suppressed during learning relative to baseline (p=0.046; Tukey’s HSD), resulting in a decreased rate relative to controls (p=0.016). Rate increases in the two weeks post-learning (p=0.0005), resulting in a higher rate than controls (p=0.05). **f,** Rehearsal modulates oligodendrogenesis rate (F_2,14_=10.33, p=0.002). Rate decreases between two weeks post-learning and rehearsal (p=0.0009), but does not differ between untrained and rehearsal mice. **g,** Consistent changes in oligodendrogenesis rate (both the increase two weeks post-learning and decrease during rehearsal) are restricted to layer I of cortex (p=0.037 and p=0.027, respectively). **h, i,** Learning modulates pre-existing sheath stability (F_5,42_=69.72, p<0.0001). Learning mice have fewer stable sheaths (p<0.0001) and more retracting sheaths (p=0.014; sheaths pseudocolored in h). **j,** Learning modulates distance of sheath change (F_2,315_=4.78, p=0.009). Sheaths retract further in learning vs untrained mice (p=0.0021). **k,**“Growing” sheaths lengthen before learning (Wilcoxon Rank-Sum; p<0.0001) but cease growth after the onset of learning. “Retracting” sheaths are initially stable but retract during (p=0.002) and after learning (p<0.0001). *\*p*<*0.05*, *\*\*p*<*0.01*, *\*\*\*p*<*0.001*.

To separate the effects of motor skill learning from motor performance, we performed *in vivo* imaging during initial training in the task (“learning”; 7 consecutive days of 20 minute reaching sessions), or during performance of the task one month following initial training (“rehearsal”; 5 consecutive days of 20 minute reaching sessions for 3 weeks; **Fig. 1b**). Of all trained mice, 93% were able to learn the task, and both learning and rehearsal of the forelimb reach task resulted in skill refinement (**Supplementary Fig. 1**). We trained mice between 2-3 months of age, when oligodendrogenesis is ongoing (**Supplementary Fig. 4, Supplementary Video 3**).

We found that learning the reach task transiently decreased and subsequently increased the rate of oligodendrogenesis in the forelimb region of motor cortex (**Fig. 1d,e**). During learning, rate of oligodendrogenesis decreased by approximately 75% relative to age-matched controls (0.14±0.03% vs. 0.58±0.08%, respectively; rate refers to the % increase in cells over the number of days elapsed). Suppression of oligodendrogenesis was restricted to the training period and was not mediated by the effects of either handling (all mice were handled equally) or training-related diet restriction (**Supplementary Fig. 4**). Immediately following learning, the generation of new oligodendrocytes increased resulting in an almost two-fold greater rate of oligodendrogenesis relative to untrained controls (0.77±0.19% vs. 0.40±0.04, respectively; **Fig. 1e**), and remained elevated up to 3 weeks following learning (**Supplementary Fig. 4**). Performance during learning predicted the magnitude of oligodendrogenesis rate increase following learning, suggesting that more successful mice showed a greater burst in oligodendrogenesis (**Supplementary Fig. 4**). In contrast to learning, rehearsal of the skilled reach task did not alter the rate of oligodendrogenesis relative to controls (**Fig. 1f**). However, the post-learning burst in oligodendrogenesis tapered off between two weeks post-learning and the rehearsal period. Overall, mice that had been trained (both “learning” and “rehearsal”) showed higher maximum rates of oligodendrogenesis across the entire experiment relative to untrained mice (**Supplementary Fig. 4**). Previous studies show that motor skill acquisition strengthens horizontal connections between neurons in motor cortex^23^. Similarly, only layer I of the forelimb region of motor cortex demonstrated consistent changes in the rate of oligodendrogenesis after learning (**Fig. 1g**). Specifically in layer I, oligodendrogenesis rate increased approximately 75% in the two weeks following learning, and then decreased by approximately 75% between two weeks post-learning and rehearsal. Rates of oligodendrogenesis in layers II/III were variable across mice.

Next, we examined the remodeling of myelin sheaths on pre-existing mature oligodendrocytes in the context of learning. Under normal physiological conditions in young adult mice, we found a small number of myelin sheaths exhibited dynamic length changes, similar to previous descriptions in mice of similar ages^24^ (14.7±1.71%; **Fig. 1h,i; Supplementary Video 4,5**). One week following forelimb reach training, the number of pre-existing myelin sheaths that underwent remodeling was increased in learning mice relative to controls (43.46±7.82% vs. 14.74±1.71%), with the most pronounced effect on sheath retraction. Learning increased both the proportion of sheaths that underwent retraction (**Fig. 1i**) and the distance these sheaths retracted compared to untrained mice (**Fig. 1j**). Learning also modulated the timing of sheath growth and retraction; growing sheaths ceased to lengthen at the onset of learning, while learning induced the retraction of previously stable sheaths (**Fig. 1k**). We found no evidence that new myelin sheaths were generated by pre-existing oligodendrocytes in untrained or learning mice.

To further characterize how motor skill learning modulates the generation of new mature oligodendrocytes, we used longitudinal *in vivo* two-photon imaging of transgenic mice that express membrane-anchored EGFP in oligodendrocyte precursor cells (OPCs; *NG2-mEGFP*^25^). We tracked OPC migration, proliferation, differentiation, and death in contralateral forelimb motor cortex over 5 weeks, beginning 1 week prior to forelimb reach training (**Fig. 2a,b**). Similar to the effects on oligodendrogenesis, learning induced a two-fold increase in the rate of OPC differentiation in the week immediately following acquisition of the task (**Fig. 2c**, 0.59±0.10% during learning vs. 1.23±0.19% post-learning), yet OPC differentiation rate was unaffected during forelimb reach training. Neither the proliferation rate nor the rate of OPC death differed significantly across the five weeks examined (**Fig. 2d,e**). However, 5/5 mice displayed a reduction in proliferation rate (~50%) during learning relative to baseline (**Supplementary Fig. 5**). Our previous work indicated that a majority of adult OPC differentiation events occur via direct differentiation rather than through asymmetric cell division^25^. In line with these findings, we found that only 10.91±3.77% of OPCs that underwent differentiation had previously proliferated during the 5 weeks of observation. The proportion of asymmetric differentiation events was unaffected by motor learning (**Fig. 2a,f,g**). To assess whether the increase in OPC differentiation following motor learning was due to parenchymal OPCs or precursors recruited from nearby brain regions or germinal zones, we tracked the cellular fate of OPCs that migrated into or out of the imaging volume (**Supplementary Fig. 5**). Migration in or out of the imaging volume was a relatively rare event, with 4.68±0.96% of the baseline number of OPCs migrating in and 2.73±0.36% migrating out. Only 3.57±1.49% of the total proliferation events and 0.70±0.30% of the total differentiation events occurred in cells that migrated into the imaging volume. There was no effect of learning on the rate of migration into or out of the imaging volume, indicating that parenchymal OPCs residing in forelimb motor cortex directly differentiated following acquisition of the reach task.

**Fig. 2.**
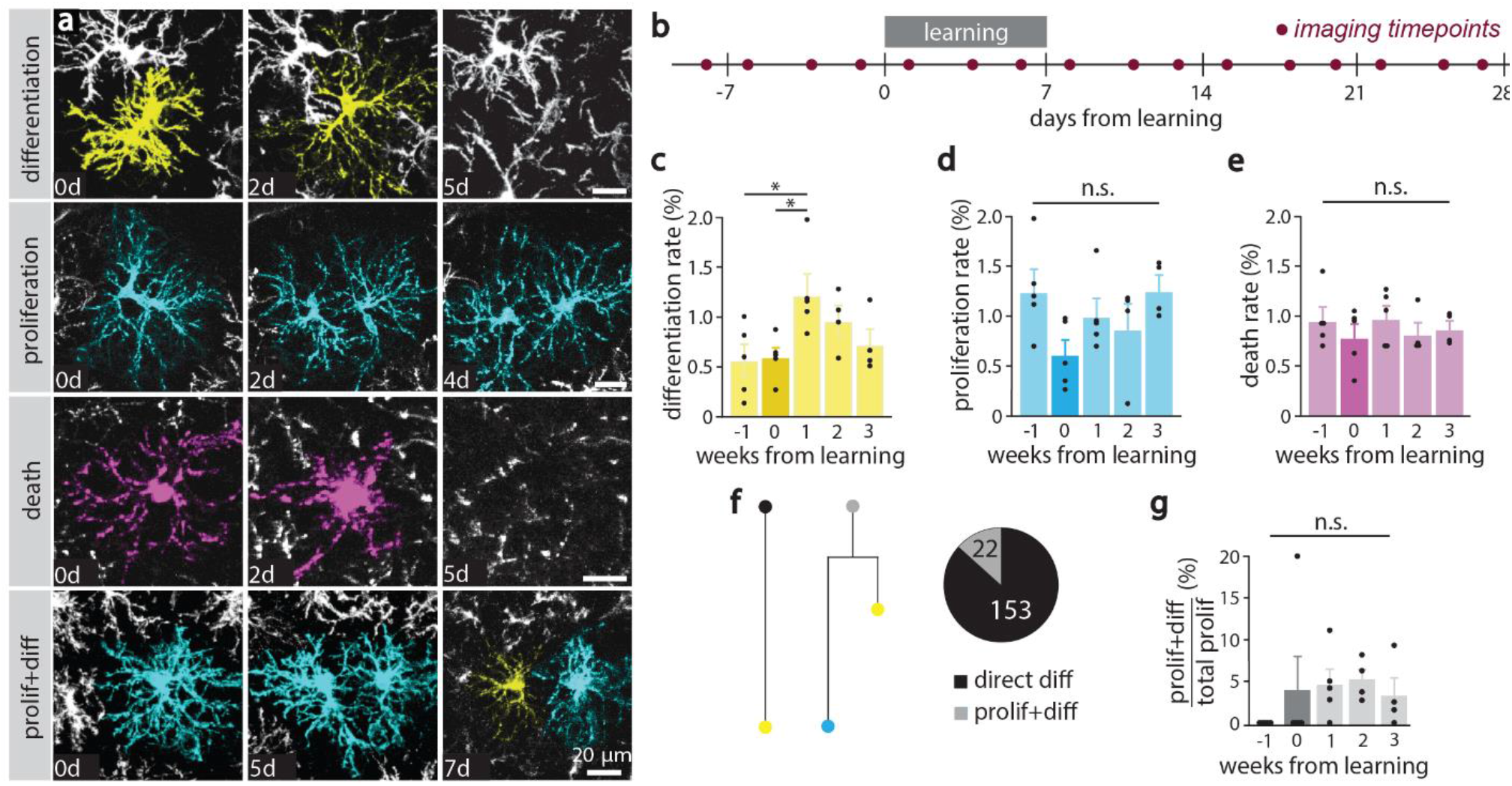
Forelimb reach learning increases OPC differentiation. **a,** *In vivo* imaging of EGFP-positive OPCs in 10 wk old *NG2-mEGFP* mice. OPCs that undergo differentiation (yellow; top) retract their filopodia, increase branching, and lose mEGFP fluorescence intensity while surrounding OPC processes infiltrate their domain. Proliferating OPCs (cyan; middle top) undergo cytokinesis and migrate to form independent domains. Dying OPCs (magenta; middle bottom) retract fragmented processes and their cell bodies become enlarged prior to disappearance. A small percentage of OPCs undergo proliferation followed by differentiation (bottom). **b,** Experimental timeline with imaging timepoints. **c,** OPC differentiation rate varies by learning week (Mean±SEM; F_4,14_=4.85, p=0.011). Rate is increased during the first week following forelimb reach training compared to both baseline and learning week (p=0.015 and p=0.021, respectively; Tukey’s HSD). **d,** No effect of learning on proliferation rate. **e,** No effect of learning on death rate. **f,** The majority (87.4%) of OPCs undergo direct differentiation (left side of cell fate diagram) as opposed to proliferation followed by differentiation (prolif+diff, right side of cell fate diagram). **g,** The proportion of differentiation events that occurred following cell division (prolif+diff) did not differ between baseline and learning or post-learning timepoints. *\*p*<*0.05*, *\*\*p*<*0.01*, *\*\*\*p*<*0.001*.

### Demyelination results in incomplete oligodendrocyte replacement, changes to the pattern of myelination, and functional deficits in motor cortex

Gray matter lesions in patients with MS contain both dying and newly forming oligodendrocytes^17^, a feature that has complicated the interpretation of remyelination in humans and animal models of MS^10,16,26^. We used longitudinal two-photon *in vivo* imaging during cuprizone-mediated demyelination to visualize the dynamics of myelin loss and repair (**Fig. 3a; Supplementary Video 6**). We fed 10-week-old congenic *MOBP-EGFP* mice a 0.2% cuprizone diet for three weeks to induce oligodendrocyte death (~90% in forelimb motor cortex, **Fig. 3a-h**), and confirmed using both SCoRe and immunohistochemistry that *in vivo* two-photon analysis of demyelination in *MOBP-EGFP* mice is a reliable measure of oligodendrocyte and myelin sheath loss (**Supplementary Fig. 6**). In contrast to the striking loss of myelin and mature oligodendrocytes, the number of cortical OPCs was unchanged following cuprizone administration relative to age-matched controls (**Supplementary Fig. 6**, 138.63±19.75 cells/mm^2^ vs. 179.11±14.99 cells/mm^2^, respectively), similar to recently published data in the cingulate cortex and the stratum lacunosum moleculare of the hippocampus^27^.

**Fig. 3.**
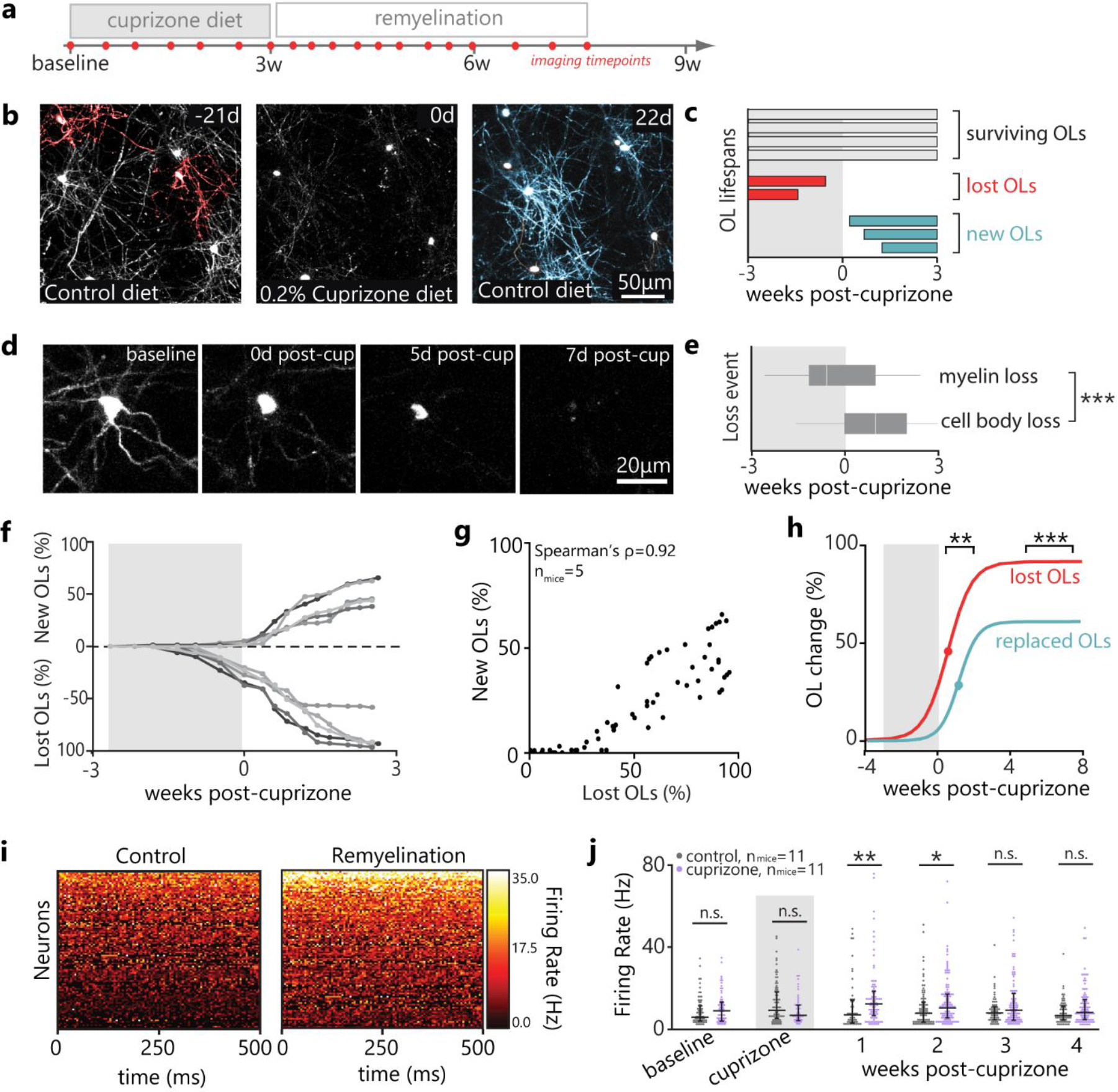
Demyelination results in incomplete oligodendrocyte replacement and functional deficits in motor cortex. **a,** Timeline of cuprizone administration and *in vivo* two-photon imaging. **b, c,** Categorization of oligodendrocyte fates following cuprizone administration as “surviving” (grey), “lost” (red), and “new” (blue). Shaded area represents cuprizone administration. **d, e,** Biphasic oligodendrocyte loss: initial loss of EGFP+ myelin sheaths and subsequent shrinking of cell body before loss of EGFP signal in an *MOBP-EGFP* mouse. e, Myelin loss (n_mice_=3, n_cells_=45) occurs earlier than oligodendrocyte soma loss (n_mice_=3, n_cells_=47; Student’s t-test, t(90)=−5, p<0.0001). **f,** Cumulative oligodendrocyte gain and loss relative to baseline (%); shades represent individual mice. **g,** Cumulative oligodendrocyte loss is tightly related to oligodendrocyte gain (Spearman’s ρ=0.922, p <0.0001). **h,** Delayed inflection point for oligodendrocyte replacement relative to loss (8.71±0.72 vs. 4.51±0.68 days post cuprizone, respectively; n_mice_=5; t(8)=4.24, p=0.0028; Student’s t-test), and decreased asymptote of replacement relative to loss (60.52±3.03% vs. 87.06±3.10%, respectively; t(8)=6.12, p=0.0003). **i,** Representative heat maps of neuronal firing rate (FR) in the motor cortex of healthy mice (left, control) versus remyelinating mice (right, cuprizone). **j,** Neuronal FR was comparable between control and cuprizone mice both prior to and during cuprizone administration, but was elevated in cuprizone mice both in the first and second week following cuprizone cessation (Wilcoxon Rank-Sum; p=0.0063 and p=0.0157, respectively). By three weeks post-cuprizone, FR was indistinguishable between cuprizone and control mice. Individual points represent individual neurons. *\*p*<*0.05*, *\*\*p*<*0.01*, *\*\*\*p*<*0.001*.

Oligodendrocyte loss occurred evenly cross cortical depths (**Supplementary Fig. 7**), leaving a small number of oligodendrocytes and myelin intact (12.94±3.10%, “surviving” cells; **Fig. 3b,c**). Cuprizone treatment suppressed oligodendrogenesis, and 85% of the few cells generated during cuprizone administration died within 3 weeks (**Supplementary Fig. 7**). Cuprizone-mediated demyelination is thought to occur through a “dying-back” process, with earliest changes in distal processes prior to cell body degeneration and demyelination^28^. Similarly, we found that oligodendrocyte death followed a biphasic model of initial myelin loss (1.53±1.27 days before the end of cuprizone) followed by cell body loss (7.26±1.22 days post-cuprizone; **Fig. 3d,e; Supplementary Video 7**). Oligodendrocyte loss plateaued approximately 15 days following the cessation of cuprizone administration (**Fig. 3f**), a conservative time course given that we designated oligodendrocyte loss as loss of EGFP fluorescence from the cell body: the last phase of oligodendrocyte death. Removal of the cuprizone diet induced a robust oligodendrogenesis response that was proportional to the extent of oligodendrocyte loss, and that plateaued after approximately three weeks (**Fig. 3f,g; Supplementary Fig. 7**). The distribution of newly generated oligodendrocytes across cortical layers following demyelination was comparable to that of oligodendrogenesis in normal conditions (**Supplementary Fig. 7**). Maximum rates of oligodendrogenesis during remyelination were 6 times greater than in healthy untrained mice (5.95±0.38% vs. 0.99±0.69%) and almost 4 times greater than in healthy trained mice (5.95±0.38% vs. 1.60±0.64%; **Supplementary Fig. 7**).

To characterize the full extent of the oligodendrogenesis response, we tracked mice up to 60 days after the end of cuprizone administration. Due to the oral uptake of cuprizone, there was inter-mouse variation in the extent of demyelination, and consequently, the extent of remyelination (**Supplementary Fig. 7**). We therefore quantified oligodendrocytes generated during remyelination as a proportion of total oligodendrocyte loss (“oligodendrocyte replacement,” %; **Supplementary Fig. 7**; See **Methods**). Oligodendrocyte replacement after cuprizone cessation followed a sigmoidal pattern, and we quantified it using three-parameter (3P) logistic equations. We were specifically interested in the inflection point (the point in time at which oligodendrogenesis switches from accelerating to decelerating) and the asymptote of the curve (the total oligodendrocyte replacement when oligodendrogenesis plateaus; **Supplementary Table 2**). Oligodendrocyte replacement was delayed relative to oligodendrocyte loss by approximately four days and plateaued significantly lower than the amount of loss (**Fig. 3h**). In fact, mice only replaced on average at 60.52±3.03% of lost oligodendrocytes in the seven weeks following removal of cuprizone diet, indicating that the remyelination response failed to completely restore baseline oligodendrocyte numbers.

To determine the effects of oligodendrocyte loss and incomplete replacement on neuronal function, we then performed chronic weekly *in vivo* extracellular recordings in the forelimb region of motor cortex in both cuprizone-demyelinated and age-matched control mice at baseline, throughout the three weeks of cuprizone treatment, and during four weeks post-cuprizone. *In vivo* multi-site electrodes record extracellular potentials from neurons within approximately a 150 micron radius of the recording electrode^29^. In healthy mice, we found that 88.57±2.36% of axon initial segments of neurons within this radius were abutted with a myelin sheath in layers II/III and V (beta-IV spectrin and MBP immunostaining along neurofilament-NeuN labeled neurons; **Supplementary Fig. 7**). In line with our *in vivo* imaging and 3P logistic models, the proportion of proximally myelinated neurons in the sampling radius was decreased by over 50% at the cessation of cuprizone treatment (38.90±3.72%), confirming we were recording from demyelinated neuronal populations. Consistent with previous work showing cuprizone application in ex vivo cortical slices does not alter synaptic transmission^30^, neuronal firing rates did not differ between groups prior to or during cuprizone administration. However, median neuronal firing rates were increased in demyelinated mice versus controls by about 70% in the first week and 40% in the second week following cuprizone cessation (11.90 vs. 6.92 Hz and 10.68 vs. 7.69 Hz, respectively; **Fig. 3i, j**), indicating they were hyperexcitable in a manner that temporally correlates with maximum oligodendrocyte loss. By three and four weeks following the removal of cuprizone diet – the time at which remyelination plateaued – neuronal firing rates in cuprizone-demyelinated mice were indistinguishable from age-matched controls. Taken together, these results demonstrate that cuprizone-mediated demyelination induces aberrant neuronal function in the forelimb region of motor cortex that recovers alongside remyelination.

Given that remyelination failed to completely restore baseline oligodendrocyte number but seemed to restore neuronal function, we sought to examine the number, length, and location of sheaths generated by new oligodendrocytes during remyelination. In the first week of remyelination, new oligodendrocytes formed more myelin sheaths compared to oligodendrocytes generated in the second week of remyelination or in control mice (54.4±3.25 vs. 39.4±1.72 and 42.28±1.30 total sheaths, respectively; **Fig. 4a-c**; **Supplementary Videos 8,9**). Similar to newborn sheaths in control mice, new sheaths generated by new oligodendrocytes during remyelination primarily grew in the first three days after generation (**Fig. 4d**) and stabilized to similar lengths as sheaths in control mice (**Fig. 4e,f**). Therefore, the increased sheath number on newborn oligodendrocytes in the week following demyelination resulted in a larger total amount of myelin generated by these oligodendrocytes (**Fig. 4g**). To examine whether oligodendrocytes generated following demyelination restored or modified the previous pattern of myelination, we examined the location of myelin sheath deposition. We found that myelin sheaths of newly generated oligodendrocytes were more often placed in previously unmyelinated areas (“remodeling”; 67.7±3.56% of sheaths) rather than in denuded areas (“remyelinating”; 32.0±3.47% of sheaths), generating a novel pattern of myelination following demyelinating injury (**Fig. 4h,i; Supplementary Videos 10,11**). These findings indicate that the myelinating capacity of individual oligodendrocytes is modulated during the phases of remyelination and that remyelination by newly generated oligodendrocytes alters the pattern of myelin in the cortex.

**Fig. 4.**
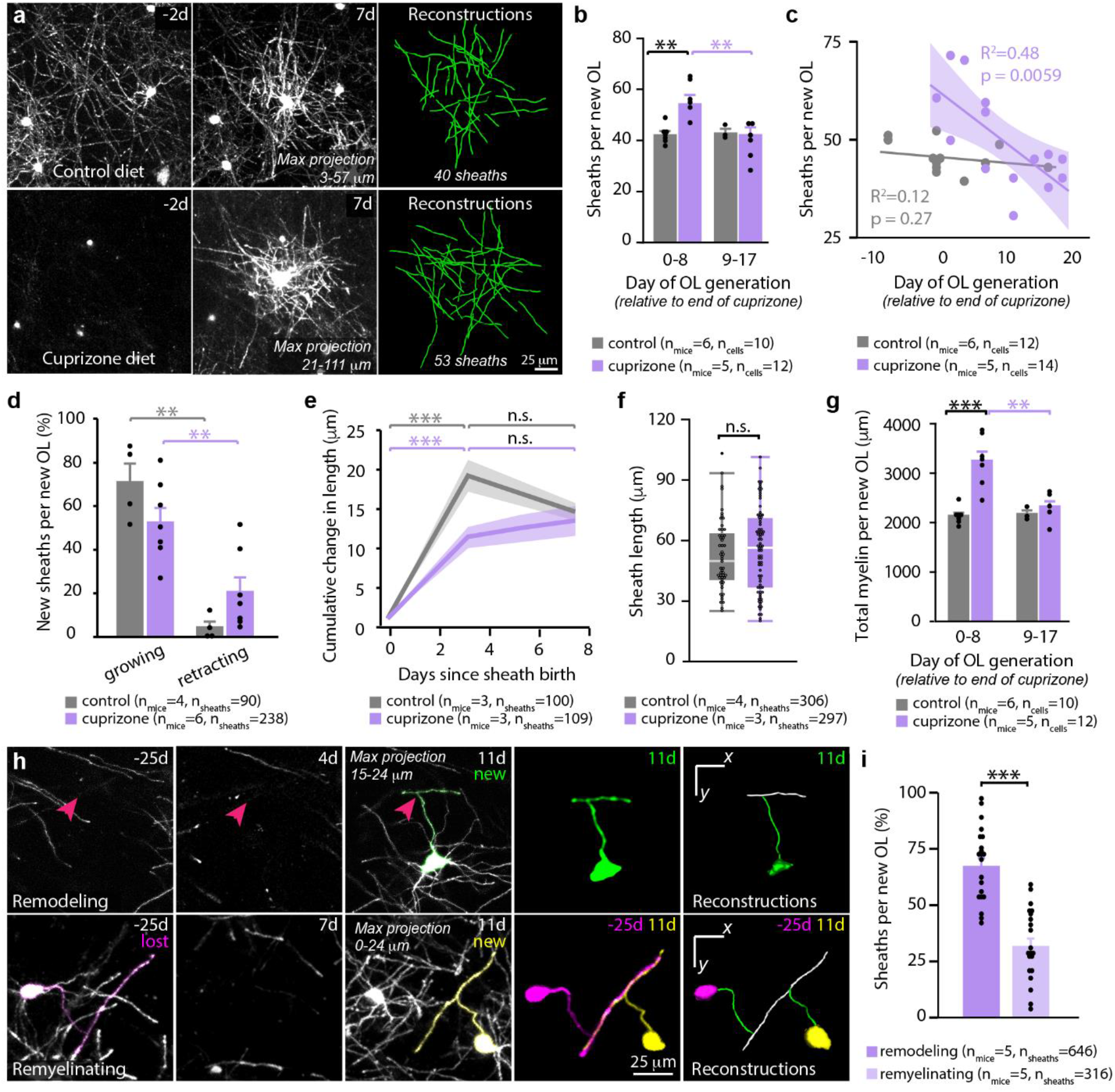
Myelin sheath number on newborn oligodendrocytes is regulated during remyelination. **a, b,** Remyelination modulates sheath number (F_1,19_=8.03, p=0.0105). Oligodendrocytes born in the first week of remyelination (top, a) generate more sheaths than age-matched control oligodendrocytes (bottom, a; p=0.010, Tukey’s HSD) and than oligodendrocytes born after week 1 of remyelination (p=0.0023). FOV shown 2 days before oligodendrocyte birth and 7 days post-birth. Points represent individual oligodendrocytes. **c,** Day of oligodendrocyte generation relative to end of cuprizone predicts sheath number in demyelinated mice (R^2^=0.48, F_1,12_=11.16, p=0.006). **d,** In the first three days post-generation, sheaths from newborn oligodendrocytes grow more often than they retract (F_3,18_=15.34, p<0.0001) in both control (p=0.0001) and cuprizone-treated mice (p=0.0096, Tukey’s HSD). Points represent individual oligodendrocytes. **e,** New sheaths change in length in the week following their generation (control: F_3,302_=47.94, p<0.0001, cuprizone: F_3,293_=29.71, p<0.0001). Sheaths in both control and cuprizone treatment stabilize their length within 3 days of sheath birth (d0 vs. d3, p<0.0001 in both treatments; Tukey’s HSD). **f,** Sheath length does not differ in control and cuprizone-treated mice 3 days after sheath generation. **g,** Remyelination shapes predicted total myelin ([mean sheath length] x [# of sheaths/OL]) generated by a newborn oligodendrocyte (F_1,19_=8.93,p=0.0077). It is higher in week 1 of remyelination than age-matched control oligodendrocytes (p<0.0001) and than oligodendrocytes born after week 1 of remyelination (p=0.0016). **h,** Newborn oligodendrocytes can place sheaths in previously unmyelinated areas (top, “Remodeling”) or previously myelinated areas (bottom, “Remyelinating”). Pink arrows point to location of junction between new sheath and new OL process. Relevant sheaths pseudo-colored. **i,** Newborn oligodendrocytes engage in remodeling more often than remyelinating (t(20)=−5.08, p<0.0001; Paired student’s t-test). *\*p*<0.05, *\*\*p*<0.01, *\*\*\*p*<*0.001*.

### Motor learning modulates oligodendrogenesis after demyelination in a timing-dependent manner

Since we found that motor learning increased both OPC differentiation and oligodendrogenesis in healthy mice (**Figs. 1,2**), we sought to examine whether motor learning could stimulate oligodendrocyte replacement in demyelinated mice. We allotted mice to one of three experimental groups: “no activity,” “early-learning” (starting 3 days post-cuprizone), and “delayed-learning” (starting 10 days post-cuprizone; **Fig. 5a, b**). Behavioral intervention had no effect on the severity of demyelination (**Fig. 5c**) nor the maximum rate of oligodendrogenesis during remyelination (**Fig. 5d**).

**Fig. 5.**
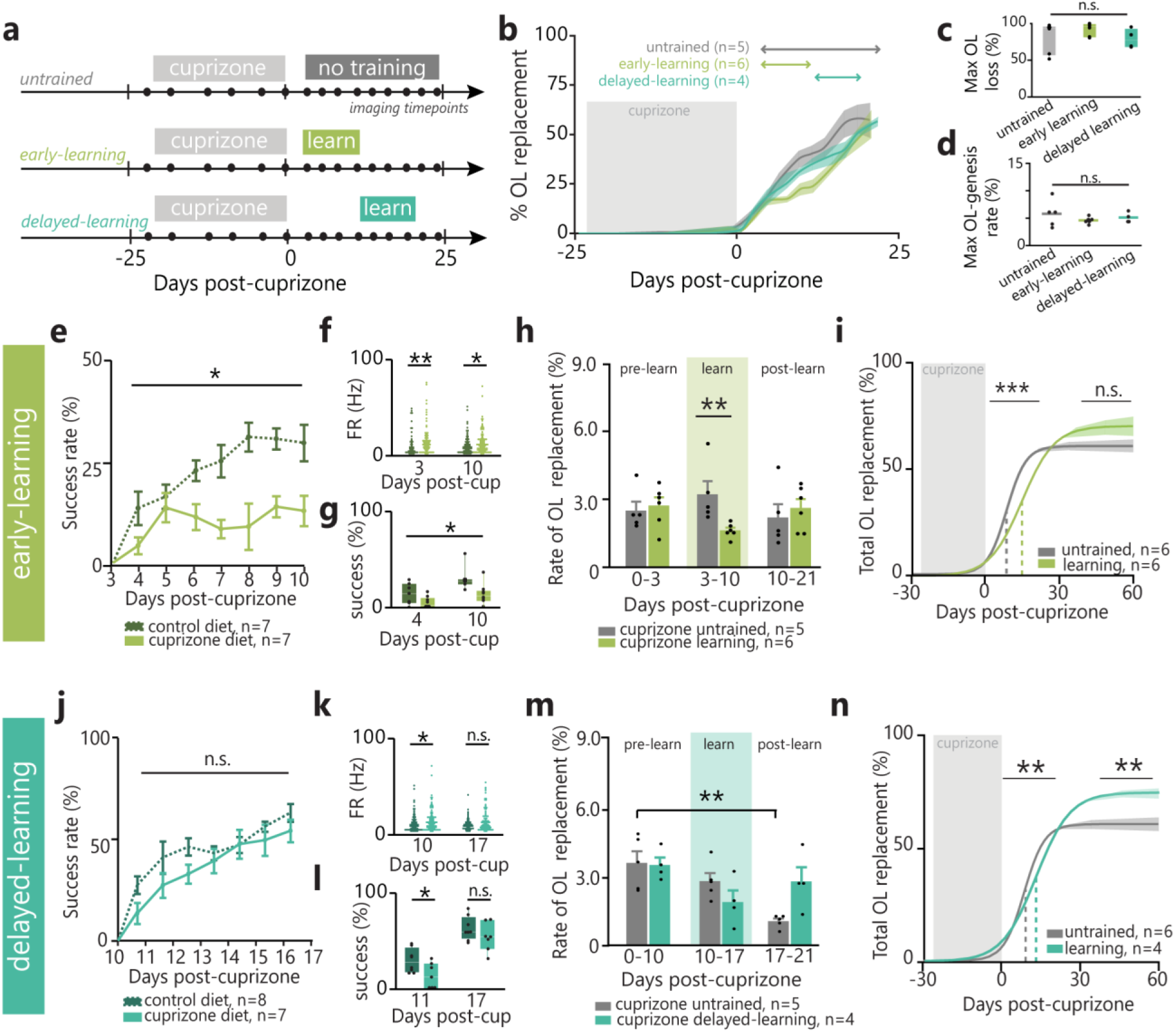
Motor learning modulates oligodendrogenesis after demyelination in a timing-dependent manner. **a,b,** Cumulative OL replacement (%; mean±SEM) across post-cuprizone behavioral interventions. **c, d,** Neither maximum OL loss nor maximum rate of oligodendrogenesis differ between behavioral interventions. **e,** Demyelination modulates early-learning success rate (F_6,78_=3.00, p=0.011). Success rate improves from first to last day of reaching for control (p=0.005), but not cuprizone-treated, mice. **f,** Both 3 and 10 days post-cuprizone, demyelinated mice have increased neuronal FR relative to controls (Wilcoxon Rank-Sum, p=0.006 and p=0.016, respectively). **g,** Both 4d post-cuprizone and 10d post-cuprizone, demyelinated mice have decreased success rates relative to controls (F_1,13_=9.09, p=0.01). **h,** OL replacement rate is suppressed during early-learning relative to untrained demyelinated mice (Wilcoxon Rank-Sum, p=0.0043). i, Delayed inflection point of OL-replacement in early-learning vs. untrained demyelinated mice (t(10)=5.77, p=0.0002), colored line/shaded area represents asymptote±SEM. **j,** No effect of cuprizone treatment on overall delayed-learning performance. **k,** 10 days post-cuprizone, but not 17 days post-cuprizone, demyelinated mice have increased neuronal FR relative to controls (p=0.016). **l,** 11 days post-cuprizone, but not 17 days post-cuprizone, demyelinated mice have impaired reaching performance relative to controls (t(12.28=−2.39, p=0.033). **m,** Delayed-learning modulates OL replacement rate (F_2,14_=4.61). Rate decreases in untrained, but not delayed-learning, mice by 21 days post-cuprizone (p=0.008). **n,** Delayed inflection point (t(8)=4.33, p=0.0025) and increased asymptote of OL-replacement (t(8)=3.35, p=0.01) in delayed-learning vs. untrained mice. *\*p*<*0.05*, *\*\*p*<*0.01*, *\*\*\*p*<*0.001*.

Mice trained in the forelimb reach task 3 days post-cuprizone (“early-learning” group) showed significant performance impairments relative to healthy controls and did not improve their reaching across the learning period, indicating a failure to acquire the reach task (**Fig. 5e; Supplementary Fig. 8**). While cuprizone did not alter overall reach attempts, the extent of demyelination was negatively related to performance (**Supplementary Fig. 8**). Motor deficits and failure to learn the forelimb reach task temporally correlated to neuronal hyperexcitability in the forelimb region of motor cortex: firing rate was increased in demyelinated mice versus healthy controls in the 10 first days post-cuprizone, coinciding with both the onset and end of the early-learning period (**Fig. 5f,g**). Learning suppressed the rate of oligodendrocyte replacement by approximately 50% in early-learning versus untrained remyelinating mice (1.62±0.13% vs 3.21±0.59%, respectively; **Fig 5b,h**), resulting in a delayed inflection point of oligodendrocyte replacement (15 days vs. 9 days post-cuprizone, respectively; **Fig. 5i**). However, the asymptote of post-cuprizone oligodendrogenesis did not differ between untrained and early-learning mice, indicating that there were no long-term effects of suppression on the extent of oligodendrocyte replacement. The learning-induced suppression was less severe in remyelinating mice than in healthy controls (50% vs. 75%, respectively; see **Fig. 1**) and we did not find an increase in the rate of oligodendrogenesis following training (**Fig. 5h**). Success rate during learning was unrelated to the asymptote of oligodendrocyte replacement in individual mice (**Supplementary Fig. 8**). In sum, motor performance was impaired and motor cortex neurons were hyperexcitable following demyelination, and failing to learn the forelimb reach task provided no benefit to oligodendrogenesis during remyelination.

Given that mice trained immediately following cuprizone cessation were unable to learn the forelimb reach task, we trained mice ten days following removal of cuprizone (ie, the half-maximum of the remyelination response; **Fig. 3h**). These “delayed-learning” mice showed no overall impairments in reaching performance (**Fig. 5j**) or reaching attempts (**Supplementary Fig. 8**) relative to healthy mice. Again, we found that neuronal hyperexcitability was temporally correlated to reaching success. While demyelinated mice were slightly less successful than healthy controls on the initial day of training (10 days post-cuprizone, when demyelinated mice still show motor cortex neuronal hyperexcitability; **Fig. 5k,l**), their success rates were indistinguishable from controls by the end of training (17 days post-cuprizone, when demyelinated mice have no difference in firing rate relative to controls). Unlike mice trained immediately after cuprizone, delayed-learners demonstrated only a slight decrease in oligodendrogenesis rate during learning (~30%) that was not statistically different from untrained demyelinated mice (**Fig. 5b,m**). While the rate of oligodendrogenesis slowed significantly by three weeks post-cuprizone in untrained mice, it did not in delayed-learners. As a result, the inflection point of oligodendrocyte replacement was delayed by over four days in delayed-learning versus untrained mice (13 vs. 9 days post-cuprizone, respectively; **Fig 5n**) and oligodendrocyte replacement plateaued substantially higher than in untrained mice (74.56±2.26% vs. 60.52±3.03%, respectively). Success during delayed-learning was not related to oligodendrocyte replacement in individual mice (**Supplementary Fig. 8**). In sum, partial remyelination restored both neuronal function and the a bility to learn the forelimb reach task, and motor learning following partial remyelination promoted long-term oligodendrogenesis.

To evaluate the effects of overall motor behavior rather than motor learning, we also trained mice prior to cuprizone administration and had them rehearse the forelimb reach task post-cuprizone (**Supplementary Fig. 9**). Although demyelinated mice demonstrated performance deficits during rehearsal, rehearsal did not modulate any aspect of remyelination. Only learning the reach task (“delayed-learning”), but not attempting to learn it (“early-learning”) nor rehearsing it (“rehearsal”), promoted oligodendrogenesis post-cuprizone.

By 7 weeks post-cuprizone, mice trained in the delayed-learning context had replaced over 20% more of lost oligodendrocytes and had over 40% greater density of oligodendrocytes in layers I-III of motor cortex than age-matched, demyelinated, untrained mice (79.24±4.56% vs 58.43±5.26%, and 86.67±5.36 vs. 60.67±4.06, respectively; **Fig. 6a-c**). Delayed-learning did not modulate the number of sheaths per new oligodendrocyte (**Fig. 6d**) but did increase the proportion of sheaths that retracted over time (**Fig. 6e**), similar to observations in healthy mice (**Fig. 1i-k**). Sheaths placed by new oligodendrocytes were equally likely to remyelinate denuded axons in both untrained and delayed-learning mice (**Fig. 6f**). Using mean sheath number per new oligodendrocyte per mouse, we modelled restoration of baseline sheath number and remyelination of denuded axons to a population level. While untrained and delayed-learning mice replaced similar proportions of baseline sheath number prior to behavioral intervention (19.59±0.97% versus 24.05±6.54%, respectively; **Fig. 6g**), delayed-learners replaced almost twice as many lost sheaths as untrained mice following training (62.22±8.12% versus 34.72±8.84%) due to prolonged oligodendrogenesis (**Fig. 5n, Fig. 6a-c**). As a result, by seven weeks post-cuprizone, delayed-learners had replaced almost 90% of their baseline sheath number, while untrained mice had only replaced on average 54% (**Fig. 6h**). Given that delayed-learners generated more sheaths that were equally likely to be deposited in a previously myelinated location as in untrained mice, we predict this resulted in almost two-fold greater remyelination of denuded axons (30.19±1.33% versus 16.38±2.37%, respectively; **Fig. 6i**).

**Fig. 6.**
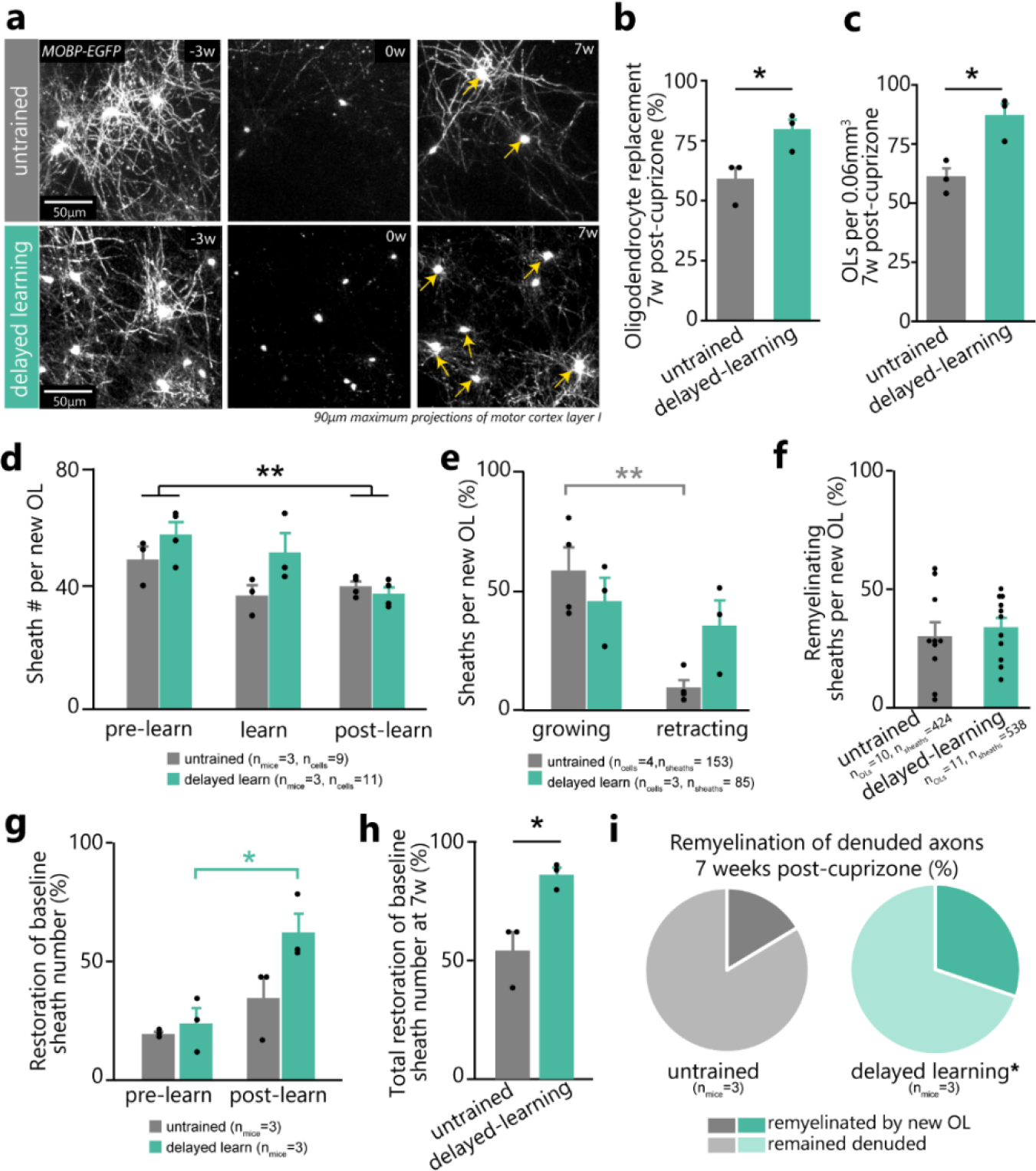
Delayed motor learning promotes remyelination via new oligodendrocytes. **a,** Representative maximum-projections of superficial cortical oligodendrocytes (OLs) at baseline (left; −3 weeks), end of cuprizone diet (middle; 0 weeks), and following 7 weeks of remyelination (right) in untrained (top) and delayed-learning (bottom) mice. Yellow arrows designate new oligodendrocytes. **b, c,** Delayed-learners replace a greater proportion of lost oligodendrocytes (t(3.92) = −2.99, p = 0.04) and have a higher density of cortical oligodendrocytes than untrained mice (t(3.72)=−3.87, p=0.02) by 7-weeks post-cuprizone (points represent individual mice). **d,** While newborn OLs have increased sheath numbers in first versus third week post-cuprizone (F_5,15_=5.14, p=0.006; p=0.0038), delayed-learning does not modulate this relationship (p=0.1; points represent individual OLs.) **e,** Delayed-learning modulates sheath dynamics (F_3,10_=6.65, p=0.0095). Sheaths on new OLs are more likely to grow than retract in untrained (p=0.007) but not delayed-learning mice (p>0.8; points represent individual OLs.) **f,** Sheaths from newborn OLs are equally likely to remyelinate denuded axons in untrained and delayed-learning mice (t(16.08)=−0.52, p=0.6; points represent individual OLs.) **g,** Population-level extrapolations suggest that delayed-learning modulates restoration of baseline sheath number (F_3,8_=7.80, p=0.0093). More sheaths are replaced after training in delayed-learning mice (Tukey’s HSD; p=0.018). **h,** Population-level extrapolations suggest that delayed-learners restore a greater proportion of baseline sheath number 7 weeks post-cuprizone (t(2.64)=−3.76, p=0.0407). **i,** Extrapolating sheath location probability to the population-level suggests that delayed-learners remyelinate a greater proportion of denuded axons than untrained mice (31% vs. 19%, respectively; t(3.14)=−5.07, p=0.013). *\*p*<*0.05*, *\*\*p*<0.01, *\*\*\*p*<*0.001*.

### Motor learning promotes the participation of pre-existing mature oligodendrocytes in remyelination

While pre-existing myelin sheaths can be remodeled under normal conditions^21,24^ (also see **Fig. 1**), current evidence suggests that mature oligodendrocytes do not participate in remyelination in rodents^5,14,31^. However, recent studies imply that myelin can be regenerated by mature oligodendrocytes in large animal models and patients with MS^10,26,32^. To determine the contribution of pre-existing mature oligodendrocytes to remyelination, we used longitudinal *in vivo* imaging and semi-automated tracing^22^ to reconstruct myelin sheaths and connecting processes to the oligodendrocyte cell body (**Supplementary Fig. 3**). We then tracked individual myelin sheaths of individual oligodendrocytes throughout cuprizone-mediated demyelination and subsequent remyelination (**Supplementary Fig. 10**; see **Methods**). Oligodendrocyte survival was variable between mice but did not differ in untrained and delayed-learning groups (12.29±7.32% vs. 20.84±6.60%; **Supplementary Fig. 11**). After three weeks of cuprizone treatment in untrained mice, all surviving oligodendrocytes experienced sheath loss, and in rare instances (1/19) oligodendrocytes added a new sheath (**Fig. 7a-d; Supplementary Videos 12,13**). While pre-existing myelin sheaths in healthy mice underwent low levels of myelin remodeling (see **Fig. 1**), cuprizone treatment increased pre-existing sheath retraction in surviving oligodendrocytes (17.0±4.22% vs. 43.8±5.95%; **Supplementary Fig. 12**). Motor skill learning one week after the removal of the cuprizone diet (delayed-learning) did not affect the degree of remodeling in pre-existing myelin sheaths during remyelination (**Supplementary Fig. 12**). In contrast, delayed-learning dramatically increased the number of pre-existing oligodendrocytes that generated new myelin sheaths (**Fig. 7d,e**). In delayed-learning mice, pre-existing oligodendrocytes were able to generate sheaths even 1.7 months after the onset of imaging, suggesting that the ability to generate myelin sheaths is an extended property of oligodendrocytes (**Fig. 7f**).

**Fig. 7.**
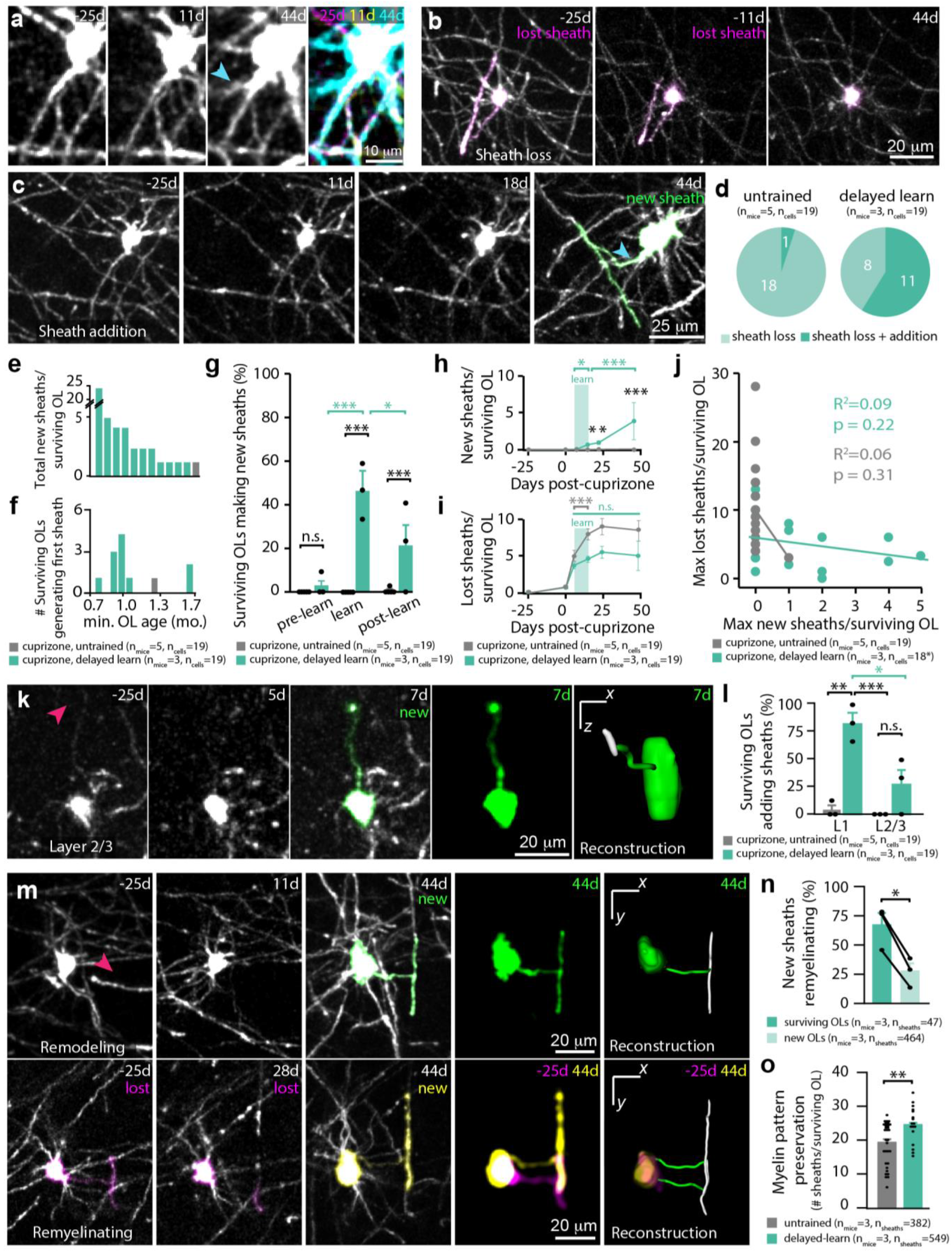
Delayed motor learning stimulates surviving mature oligodendrocytes to contribute to remyelination. **a,** Identification of surviving oligodendrocytes (OLs) via conserved processes. Note the new process pointed out on the same oligodendrocyte in **a** and **c** (cyan arrow). **b,c,d,** Surviving OLs both lose (pink) and generate sheaths (green). Image manually resliced in **c** (44d) to show sheath and process connecting to cell body. **e,f,** Number of sheaths generated per surviving OL and minimum possible OL age at time of sheath generation (assuming age 0 at imaging onset). **g,** Delayed-learning modulates surviving OL sheath production (F_2,51_=9.30, p=0.0004). Sheath generation increases during learning (p<0.0001) and decreases post-learning (p=0.019), resulting in elevated generation relative to untrained mice both during (p<0.0001) and after learning (p=0.026). **h,** Learning modulates cumulative new sheaths on surviving OLs (F_7,618_=12.96, p<0.0001). Delayed-learning increases new sheaths relative to baseline (p=0.019) and relative to untrained mice both during (p=0.028) and after (p=0.033) learning. Sheath number increases up to 4 weeks post-learning (p<0.0001). **i,** Learning modulates cumulative lost sheaths on surviving OLs (F_7,611_=7.04, p<0.0001). Sheath loss initially increases in untrained and delayed-learning mice (p<0.0001 and p<0.0001, respectively) then ceases in delayed-learning (p>0.9) but not untrained mice (p<0.0001). **j,** No relationship between sheath loss and gain (single outlier removed for analysis). **k,l,** Learning increases sheath generation by surviving OLs in both L1 and L/3 relative to controls (F_1,6_=7.05, p=0.038; p=0.0019 and p=0.0016, respectively), though generation is heightened within L1 versus L2/3 (p=0.044). **m,n,** New sheaths from surviving OLs are more likely to remyelinate denuded axons than sheaths from newborn OLs (t(2)=7.28, p=0.018). Pink arrows point to location of junction between new sheath and surviving OL process. Relevant sheaths pseudo-colored. **o,** Surviving OLs in delayed-learning mice contribute more sheaths to the original pattern of myelination (via maintenance and addition) than untrained mice (Wilcoxon Rank-sum, p=0.004). *\*p*<*0.05*, *\*\*p*<*0.01*, *\*\*\*p*<*0.001*

We found that the generation of new sheaths from pre-existing oligodendrocytes was temporally correlated with the onset of learning the forelimb reach task. During learning, the number of pre-existing oligodendrocytes generating new sheaths increased by over 40%, and pre-existing oligodendrocytes continued to generate sheaths even following learning (**Fig. 7g; Supplementary Video 13,14**). This corresponded with a significant increase in cumulative new sheaths generated by surviving oligodendrocytes in delayed-learning mice relative to untrained mice, both during and after learning (**Fig. 7h**). Myelin sheath loss stagnated in surviving oligodendrocytes after the onset of learning, in contrast to sheath loss in untrained mice which continued for two weeks following cuprizone (**Fig. 7i**). Sheath loss did not affect the number of new sheaths generated on an individual oligodendrocyte (**Fig. 7j**).

As with oligodendrogenesis following learning, we found that new sheath addition in pre-existing oligodendrocytes was higher in layer I compared to layer II/III of cortex (**Fig. 7k,l; Supplementary Video 15**). Similar to newly generated oligodendrocytes following demyelination, surviving oligodendrocytes formed new myelin sheaths on both denuded and previously unmyelinated axons (**Fig. 7m; Supplementary Videos 16,17**). When comparing remyelination by new sheaths from surviving oligodendrocytes and newly generated oligodendrocytes, a significantly larger proportion of surviving oligodendrocyte sheaths remyelinated denuded axons (**Fig. 7n**). The combination of learning-induced cessation of sheath loss and new sheath generation from surviving oligodendrocytes resulted in significantly more sheaths from surviving oligodendrocytes maintaining the original myelination pattern in delayed learning mice relative to untrained mice (**Fig 7o**).

Pre-existing oligodendrocytes engaging in new myelin sheath deposition showed an increase in overall cell body volume of 141±15% (**Supplementary Fig. 13**). Sheath generation in pre-existing oligodendrocytes followed a similar time course to sheath generation in newborn oligodendrocytes, with sheaths growing during the first three days of their generation and then stabilizing in length (**Supplementary Fig. 11**). In healthy mice, pre-existing oligodendrocytes were never observed generating new sheaths (**Supplementary Fig. 11**). These findings indicate that, following demyelination but not under normal conditions, motor learning specifically enhances the ability of pre-existing oligodendrocytes to generate additional myelin and maintain pre-existing sheaths (**Supplementary Fig. 14**).

## Discussion

Tissue regeneration following injury or disease is a long sought-after goal, particularly in the adult nervous system. Oligodendroglia represent one of the few cell types that retain the capacity to regenerate and repair following damage to the adult CNS. Remyelination of denuded axons can restore neuronal function^6^, promote neuroprotection^33^, and may facilitate functional recovery in CNS diseases characterized by myelin loss^8,9^.

In this study, we show that learning not only shapes neuronal circuits but also their pattern of myelination in both the healthy brain and during recovery from neurological injury. Longitudinal *in vivo* two-photon imaging of the forelimb motor cortex throughout the learning and rehearsal of a forelimb reach task revealed that learning transiently suppressed oligodendrogenesis but subsequently increased oligodendrocyte generation, OPC differentiation, and retraction of pre-existing myelin sheaths. Next, we examined the *in vivo* dynamics of myelin loss and repair using a cuprizone-mediated demyelination model, which induced ~90% motor cortical oligodendrocyte loss, neuronal firing rate abnormalities, and deficits in motor performance. Motor learning modulated the dynamics of remyelination in a timing-dependent manner. When reach training occurred immediately following demyelination and motor cortex neurons were hyperexcitable, mice were unable to learn the task and had no benefit to long-term recovery. However, when training occurred ten days after the onset of remyelination and neurons were no longer hyperexcitable, mice were able to learn the task, resulting in greater oligodendrocyte generation and myelin sheath replacement. In addition, motor learning enhanced the ability of surviving oligodendrocytes to participate in remyelination via the generation of new sheaths. These results demonstrate that motor learning can modulate myelin repair both via cortical oligodendrogenesis and via myelin sheath formation by surviving oligodendrocytes to improve remyelination outcomes (**Supplementary Fig. 14**).

To our knowledge, we are the first to observe a transient learning-induced suppression in oligodendrocyte generation. OPC differentiation was unaffected during this suppression (**Fig. 2**), suggesting that learning may temporarily decrease the survival and integration of differentiated OPCs as mature myelinating oligodendrocytes, in line with previous work in the developing CNS^34^. These nuanced effects may be apparent in our study due to high intra-individual resolution and regional specificity of oligodendrocyte lineage cells under observation, whereas previous studies did not parcellate motor cortex based on function for data analysis^13,18,19^. Motor learning requires precise network connectivity of both intracortical and corticospinal synapses^20^ and the consolidation of associated spatiotemporal activity patterns^35^. It is possible that location-specific cues suppress the integration of new oligodendrocytes to prevent aberrant myelination during learning, or that metabolic demand imposed by network plasticity during learning^36^ may deplete the resources required for the generation and integration of adjacent oligodendrocytes^37^. How these learning-induced changes are communicated to the oligodendrocyte lineage cells remains undefined. Axons form synapses with local OPCs, and neuronal activity can modulate OPC proliferation and differentiation within both the healthy CNS and demyelinated regions^18, 38–41^. This communication may be mediated by the effects of brain-derived neurotrophic factor (BDNF) on both activity-dependent synaptic modulation^42^ and oligodendrocyte maturation and myelination^43,44^. Future work characterizing mechanisms underlying activity-dependent oligodendrogenesis particularly during different life stages will provide additional insight on promoting recovery from demyelinating injury.

Previously, we reported that environmental enrichment of middle-aged mice leads to increased oligodendrogenesis, but not to the remodeling of myelin sheaths in the somatosensory cortex^21^. Here, we found that motor learning increased remodeling of pre-existing myelin sheaths in the motor cortex of young adult mice. This disparity may be due to age-related decreases in the ability of pre-existing myelin sheath remodeling, as is observed with synaptic plasticity^45^. Alternatively, heterogeneity^46^ or regional variance between somatosensory and motor oligodendrocytes may play a role in this differential response. The role of sheath retraction throughout learning remains to be explored, though recent work suggests this may regulate conduction velocity^47,48^ and allow for increased axonal branching^49–51^. Concurrent multi-color *in vivo* imaging of reach-specific neurons and myelin sheath dynamics in the context of learning will provide further information into how neurons and oligodendrocytes interact to facilitate the consolidation of neural circuits.

The consequences of demyelination on the function of intact cortical neural circuits is not well understood. We found that neural firing rate was increased following demyelination and resulted in hyperexcitability of the forelimb motor cortex. Despite effects of copper chelation on NMDAR and AMPAR currents, LTP suppression, and neuronal excitability^52–54^, we found no effect of cuprizone treatment on concurrent neuronal firing rate *in vivo*. This corroborates whole cell recordings of layer 5 pyramidal neurons showing that cuprizone-mediated demyelination, but not bath application of cuprizone, results in increased intrinsic excitability^30^. Hyperexcitability is a common feature of neurodegenerative diseases and can be a consequence of increased cellular stressors, including elevated intracellular calcium load^55^. Within neuroinflammatory lesions in the spinal cord caused by an experimental autoimmune encephalomyelitis (EAE) model of MS, axons show elevated calcium load, with highest levels in demyelinated axons^56^. The accumulation of cellular stress as a result of demyelination may be a potential cause of neurodegeneration seen in patients with progressive MS.

In addition to cell-autonomous mechanisms, hyperexcitability may also result from changes in excitatory and inhibitory balance within motor cortex circuits. Motor cortex control of movement results from excitatory input from other brain regions that is processed by complex local excitatory and inhibitory circuitry^57^. Hyperexcitability, and subsequent behavioral impairment, could be due to alterations in the myelination of thalamocortical input, corticospinal output axons, or of local circuitry within primary motor cortex. In neurodegenerative disorders, local circuit processing alterations produce hyperexcitability^58–60^, which predicts imminent motor impairments in both human patients and animal models^61,62,63^. We found that neuronal activity in remyelinating mice was indistinguishable from healthy controls when superficial motor cortex remyelination plateaued, and that this partial remyelination rescued motor deficits relative to healthy controls. These data show a clear link between cortical myelin content, neuronal function, and motor behavior. Future work characterizing the cell-type specific changes in neuronal excitability^64^ and plasticity^65^, along with circuit-level alterations in excitatory and inhibitory balance within motor cortex^66^ will provide insights into the cellular and circuit origins of demyelination-induced hyperexcitability.

Recent analysis of tissue from MS patients suggests that shadow plaques are incompletely demyelinated lesions or lesions remyelinated by pre-existing oligodendrocytes^10,26^. Our data clearly demonstrate both scenarios occur following demyelinating injury. Furthermore, we show that the remyelinating potential of pre-existing mature oligodendrocytes is modulated by learning-induced plasticity. While the majority of surviving oligodendrocytes contributed new myelin sheaths during remyelination in response to learning, the variability in the magnitude of the response raises questions about what drives certain oligodendrocytes to generate more new myelin than others. One possibility is that proximity to a learning-relevant neural circuit modulates the amount of new myelin deposited by surviving oligodendrocytes^67,68^. Exploring whether surviving oligodendrocytes preferentially myelinate axons that they are already ensheathing may provide insight as to whether direct axo-myelinic communication^69^ may play a role in initiating the remyelination response in certain pre-existing mature oligodendrocytes over others. The continued reduction in cell soma size and progressive compaction of chromatin of newly generated oligodendrocytes are hallmarks of the maturation of OPCs into myelinating oligodendrocytes^21,70,71^. Cell soma volume increases during remyelination suggests that epigenetic regulation of gene expression in pre-existing mature oligodendrocytes may regulate the generation and deposition of new myelin, similar to recent findings in MS patient tissue^26^. Identification of gene expression changes in mature oligodendrocytes engaging in new sheath deposition could lead to insights into the molecular mechanisms involved in remyelination by pre-existing oligodendrocytes and provide new targets for therapies to promote remyelination.

Behavioral interventions used to support motor function in MS patients^15^ may also engage endogenous mechanisms to stimulate oligodendrogenesis. We found that the learning-induced burst in oligodendrogenesis can be used to enhance remyelination after injury. Notably, motor learning implemented after the onset of remyelination prolonged the duration of oligodendrogenesis, increased long-term oligodendrocyte replacement, nearly restored baseline sheath number, and promoted the remyelination of denuded axons by both new and surviving oligodendrocytes (**Supplementary Fig. 14**). Overall, our findings argue that specifically timed therapeutic interventions designed to promote oligodendrogenesis and engage mature oligodendrocytes in myelin repair may enhance remyelination and speed functional recovery in demyelinating disorders.

## Acknowledgements

We thank Anthony Chavez for technical assistance; Andrew Scallon and the Optogenetics and Neural Engineering Core (P30NS048154); Dominik Stitch and the University of Colorado Anschutz Medical Campus Advance Light Microscopy Core for help with SCoRE imaging (P30NS048154); Mike Hall for machining expertise; Samantha Bromley-Coolidge for assistance with reach videos; S. Rock Levinson for discussions on 3P logistic equation modeling; Bob Cudmore for image processing expertise; Matthew Rasband for providing his βIV spectrin (C9581) antibody; and members of the Hughes and Welle labs for discussions.

## Funding

MAT is supported by the NIH Institutional Neuroscience Graduate Training Grant (5T32NS099042-17). Funding was provided by NIH 1R21EY029458-01 and the Boettcher Foundation Webb-Waring Biomedical Research Award to CGW and the Boettcher Foundation Webb-Waring Biomedical Research Award, the Whitehall Foundation, the Conrad N. Hilton Foundation (17324), and the National Multiple Sclerosis Society (RG-1701–26733) and NINDS (NS106432) to EGH.

## Author Contributions

EGH and CRM conceived the project. CMB designed, analyzed, and generated Figs. 1,4,6,7; Supp. Figs. 2,3,6,10-13; Supp. Videos 2-17. HJB designed, analyzed, and generated Figs. 1,3,5,6; Supp. Figs. 1,4,7-9,14, Video 1. CRM contributed to data analysis in Fig. 1,3,5, Supp. Fig. 1,7,8. MAT designed, analyzed, and generated Fig. 2; Supp. Fig. 2,5,6. DN performed all electrophysiological experiments and analyses. CGW supervised the electrophysiological experiments. EGH supervised the project. CMB, HJB, and EGH wrote the manuscript with input from other authors.

## Competing interests

The authors declare no competing financial interests.

## Data and materials availability

All data that support the findings, tools and reagents will be shared on an unrestricted basis; requests should be directed to the corresponding author.

## List of Supplementary materials

Supplementary Tables 1 and 2

Supplementary Figs. 1 to 14

Supplementary Videos 1 to 17

## Methods

### Animals

All animal experiments were conducted in accordance with protocols approved by the Animal Care and Use Committee at the University of Colorado Anschutz Medical Campus. Male and female mice used in these experiments were kept on 14h light/10h dark schedule with ad libitum access to food and water, aside from training-related food restriction (see *Forelimb Reach Training*). All mice were randomly assigned to conditions and were precisely age-matched (±5 days) across experimental groups. *NG2-mEGFP* (Jackson stock #022735) and congenic C57BL/6N *MOBP-EGFP* (MGI:4847238) lines, which have been previously described^72,73^, were used for two-photon imaging. Wild-type C57\B6N Charles River wild-type mice were used in electrophysiological experiments.

### Two-photon microscopy

Cranial windows were prepared as previously described^73^. Six- to eight-week-old mice were anesthetized with isoflurane inhalation (induction, 5%; maintenance, 1.5–2.0%, mixed with 0.5 L/min O2) and kept at 37 °C body temperature with a thermostat-controlled heating plate. After removal of the skin over the right cerebral hemisphere, the skull was cleaned and a 2 × 2 mm region of skull centered over the forelimb region of primary motor cortex (0 to 2 mm anterior to bregma and 0.5 to 2.5 mm lateral) was removed using a high-speed dental drill. A piece of cover glass (VWR, No. 1) was then placed in the craniotomy and sealed with Vetbond (3M) and subsequently dental cement (C&B Metabond). A 5mg/kg dose of Carprofen was administered subcutaneously prior to awakening and for three additional days for analgesia. For head stabilization, a custom metal plate with a central hole was attached to the skull. *In vivo* imaging sessions began 2-3 weeks post-surgery and took place 2-3 times per week (see imaging timelines in **Figs. 1–3,5**). During imaging sessions, mice were anesthetized with isoflurane and immobilized by attaching the head plate to a custom stage. For *MOBP-EGFP* experiments, images were collected using a Zeiss LSM 7MP microscope equipped with a BiG GaAsP detector using a mode-locked Ti:sapphire laser (Coherent Ultra) tuned to 920 nm. *NG2-mEGFP* mice were imaged using a Bruker Ultima Investigator microscope equipped with Hamamatsu GaAsP detectors and a mode-locked Ti:sapphire laser (Coherent Ultra) tuned to 920 nm. The average power at the sample during imaging was 5-30 mW. Vascular and cellular landmarks were used to identify the same cortical area over longitudinal imaging sessions. *MOBP-EGFP* image stacks were acquired with a Zeiss W “Plan-Apochromat” 20X/1.0 NA water immersion objective giving a volume of 425 μm × 425 μm × 336 μm (1,024 × 1,024 pixels; corresponding to layers I–III, 0-336 μm from the meninges) from the cortical surface. *NG2-EGFP* image stacks were acquired with a Nikon LWD Plan Fluorite 16X/0.8 NA water objective with a volume of 805 μm × 805 μm × 336 μm (2,048 × 2,048 pixels; corresponding to layers I–III, 0-336 μm from the meninges).

### SCoRe microscopy

Spectral confocal reflectance microscopy (SCoRe) was performed as previously described^74^. For the *MOBP-EGFP* SCoRE/two-photon validation experiments, *in vivo* image stacks were acquired on an Olympus F1000MPE upright multiphoton microscope (DIVER). Single-photon confocal microscopy was performed using 488, 543, and 633nm laser lines combined with appropriate emission filters and descanned Olympus detectors. Two-photon microscopy of *MOBP-EGFP* fluorescence was performed immediately following SCoRe imaging using a mode-locked Insight X3 laser (Spectra-Physics) tuned to 920nm and non-descanned Olympus detectors. All images were obtained using an Olympus 20X/1.0 NA water immersion objective (XLUMPLFLN20XW). SCoRe image channels were summed, registered to the two-photon data, and then analyzed for SCoRe/two-photon colocalization using ImageJ.

### Cuprizone-mediated demyelination

Cortical demyelination was induced in our congenic C57\B6N *MOBP-EGFP* mice using 0.2% Cuprizone (Sigma Chemical #C9012), stored in a glass desiccator at 4°C. Cuprizone was mixed into powdered chow (Harlan) and provided to mice in custom feeders (designed to minimize exposure to moisture) for three weeks on an ad libitum basis. Feeders were refilled every 2-3 days, and fresh cuprizone chow was prepared weekly. Cages were changed weekly to avoid build-up of cuprizone chow in bedding, and to minimize reuptake of cuprizone chow following cessation of diet via coprophagia. We used a 3-week partial cortical demyelination model (resulting in 88.3±2.9% oligodendrocyte loss in motor cortex) to allow us to track the same area of interest over time using surviving oligodendrocytes, and to investigate the behavior of surviving oligodendrocytes. Given that cuprizone was ingested on a voluntary basis, we controlled for variation in dosage in several ways. First, we weighed a subset of mice (n = 19) before and after cuprizone diet to ensure no weight loss had occurred. We found that on average, mice gained weight during cuprizone administration, confirming their consumption of the drug (Paired student’s t-test, t(18) = 2.32, p = 0.03). Second, we investigated variation in total oligodendrocyte loss. There was variation in maximum oligodendrocyte loss (50-100%), and oligodendrocyte loss and gain had a partially homeostatic relationship in that the amount of loss significantly predicted the subsequent amount of oligodendrocyte gain (**Fig. 3g**; **Supplementary Fig. 7**). To control for variation in total oligodendrocyte loss, and its subsequent effects on oligodendrocyte gain, we therefore measured oligodendrocyte gain relative to the severity of loss using the following equation:

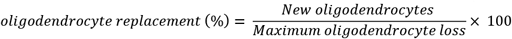

### Forelimb reach training

Mice were weighed, habituated to a training box for 20 minutes, and deprived of food 24 hours prior to training. The training box was fitted with a window providing access to a pellet located on a shelf 1cm anterior and 1mm lateral to the right-hand side of the window. After one session of initial habituation, training sessions began daily for 20 minutes. Mice learned to reach for the pellet using their left hand. Successes were counted when the mouse successfully grabbed the pellet and transported it inside the box. Errors were qualified in three ways: “Reach error” (the mouse extends its paw out the window but does not grab the pellet), “Grasp error” (the mouse reaches the pellet but does not successfully grasp onto it), and “Retrieval error” (the mouse grasps the pellet but does not succeed in returning it to the box; **Supplementary Video 1**). Mice were kept on a restricted diet throughout training to maintain food motivation but were weighed daily to ensure weight loss did not exceed 10%. For forelimb reach training, mice underwent habituation (average of ~2 days of exposure) followed by training until seven consecutive days of training with reach attempts were recorded. For the rehearsal of the forelimb reach task, mice performed the reach task in a 20-minute session, 5 days per week over three weeks. Similar to previously published findings, over 90% of mice trained in forelimb reach context were able to learn the task^75^; mice were excluded if they were not able on a single day to succeed in at least 10% of reaches (**Supplementary Fig. 1**). To control for any batch or experimenter effects in forelimb reach training results, behavioral performance was only compared for mice trained by the same experimenter within the same batch (i.e., control and experimental mice were only compared if trained at the same time by the same experimenter).

### Immunohistochemistry

Mice were anesthetized with an intraperitoneal injection of sodium pentobarbital (100 mg/kg b.w.) and transcardially perfused with 4% paraformaldehyde in 0.1 M phosphate buffer (pH 7.0-7.4). Brains were postfixed in 4% PFA for 1 h at 4 °C, transferred to 30% sucrose solution in PBS (pH 7.4), and stored at 4 °C for at least 24 h. Brains were extracted, frozen in TissuePlus O.C.T, and sectioned coronally or axially (bregma 0 to 2 mm) at 50 μm thick. Immunostaining was performed on free-floating sections. Sections were pre-incubated in blocking solution (5% normal donkey serum, 2% bovine g-globulin, 0.3% Triton X-100 in PBS, pH 7.4) for 1-4 h at room temperature, then incubated overnight at 4 °C in primary antibody (listed along with secondary antibodies in Supplementary Table 1). Secondary antibody incubation was performed at room temperature for 2 h. Sections were mounted on slides with Vectashield antifade reagent (Vector Laboratories). Images were acquired with a laser-scanning confocal microscope (Zeiss LSM 510).

### Image Processing and Analysis

Image stacks and time-series were analyzed using FIJI/ImageJ. All analysis was performed on unprocessed images except in surviving oligodendrocyte myelin sheath analysis images, which were pre-processed by with a Gaussian blur filter (radius = 1 pixels) to denoise. When generating figures, image brightness and contrast levels were adjusted for clarity. For the pseudocolor display of individual myelin sheaths or OPCs (**Figs. 1–4,6,7; Supplementary Fig. 2,3,11,12; Supplementary Videos 2,3,5,6,8-17**), a max projection of the region of interest was generated and was manually segmented and colorized. Longitudinal image stacks were registered using FIJI plugins ‘Correct 3D drift’67 or ‘PoorMan3DReg’. When possible, blinding to experimental condition was used in analyzing image stacks from two-photon imaging. To ensure the validity of oligodendrocyte lineage cell tracking, we performed interrater reliability on a subset of images and found a highly significant correlation between raters (R^2^ = 0.998, p < 0.0001).

#### Cell Tracking

Custom FIJI scripts were written to follow oligodendrocytes in four dimensions by defining EGFP+ cell bodies at each timepoint, recording xyz coordinates, and defining cellular behavior (new, dying, proliferating, differentiating, or stable cells). Mature oligodendrocyte and OPC migration, proliferation, death, and differentiation were defined as previously described^72,73^. Differentiation events were recorded as the time point immediately preceding the total loss of *NG2-mEGFP* fluorescence. OPC and oligodendrocyte gain and loss were quantified cumulatively relative to baseline cell number to account for variation in starting cell number. Rate of oligodendrocyte gain was quantified as the percent change in gain over the amount of time elapsed.

#### Identification of surviving oligodendrocytes

A subset of surviving oligodendrocytes exhibited drastic changes in morphology during remyelination in the form of cell body expansion, sheath addition, and increase in EGFP expression (**Supplementary Figs. 11-13**). To ensure that these oligodendrocytes were in fact individual surviving oligodendrocytes and not new oligodendrocytes generated in a similar location that myelinated similar axonal locations, we employed several criteria for inclusion in our final dataset. Due to the stringency and conservative nature of these criteria, we consider our findings to likely be an underestimate of the capacity of surviving oligodendrocytes to generate new myelin sheaths:

1. Change in cell soma centroid (<2.5 standard deviations from the mean)
2. Percentage of sheath retention (>10% of original sheaths)
3. Change cell soma volume (<700 μm^3^)
4. Protracted sheath addition (>6 days)
5. Semi-automated tracing of new sheath to OL surviving cell body (using Simple Neurite Tracer)
6. Distance between surviving and newborn oligodendrocyte cell bodies at time point of sheath generation (>50 μm)

We measured the change in centroid position of surviving oligodendrocyte cell bodies from baseline to day of peak remodeling—i.e. the day where the largest number of sheaths were added by a given oligodendrocyte. We ensured that the 3D location of individual surviving oligodendrocytes did not change by more than 2.5 standard deviations from the mean displacement of reference objects measured within the same stack (5.459±2.373 μm; **Supplementary Fig. 10**). Since myelin sheath loss occurs significantly earlier than cell body loss in oligodendrocyte cell death (**Fig. 3d,e**) and a new oligodendrocyte could be born in the same location immediately following the death of the original oligodendrocyte, we only included surviving oligodendrocytes that conserved at least 10% of their original processes (mean percent conserved processes = 75.6±4.45%). There is an increase in the frequency of pairs and rows of oligodendrocytes with adjacent cell somas in the aging brain^76^. Since the axial resolution of our two-photon microscope is ~2.6 μm, we should be able to resolve two directly adjacent oligodendrocytes in the z-direction. To ensure that this is the case, we took several additional steps to rule out the addition of a new oligodendrocyte generated immediately adjacent in the z-axis. Reduction of cell soma size is a hallmark of the maturation of OPCs into myelinating oligodendrocytes^73,77,78^; therefore, we measured the cell soma volume of newly generated oligodendrocytes (**Supplementary Fig. 10**). Next, we measured each surviving oligodendrocyte at baseline and at the day of peak remodeling to determine the change in cell soma volume. Assuming a z-distance of 0 μm between two cells, we excluded oligodendrocytes that grew more than 700 μm^3^, the volume of the smallest new oligodendrocyte measured (**Supplementary Fig. 10**). Additionally, we ensured that this growth was not unidirectional (as you would see with the addition of an adjacent oligodendrocyte cell body), but rather that the oligodendrocyte cell body expanded in multiple directions around the centroid position. Previous studies have indicated that new oligodendrocytes have a limited period in which to generate all of their myelin sheaths, a few hours in the developing zebrafish^79^ and less than 18 hours in vitro^80^. We found that newborn oligodendrocytes generate all sheaths within 0-3 days of generation in mice (n_mice_ = 11; n_cells_ = 26). If a new oligodendrocyte was generated directly adjacent to a surviving oligodendrocyte, we would predict that all additional myelin sheaths would be added within this time frame. Therefore, we required that surviving oligodendrocytes that added more than 4 sheaths— approximately 10% of the average sheaths generated by an oligodendrocyte— must have added these new sheaths over multiple, non-consecutive imaging time points (**Supplementary Fig. 10, Supplementary Video 14**). Using these criteria, two oligodendrocytes were excluded from the final dataset as they may have represented the addition of a new oligodendrocyte in a similar location. One oligodendrocyte violated both the change in centroid requirement and protracted sheath addition requirement (change in centroid = 19.3266 μm, and 8 sheaths were added at one time point) and the other violated the terms for protracted sheath addition (over 10 sheaths added within two consecutive timepoints). Three other surviving oligodendrocytes were excluded from new sheath analysis because newborn oligodendrocytes were generated within 50 μm of the surviving oligodendrocyte cell body on the day of surviving oligodendrocyte sheath addition. The surviving oligodendrocyte dataset was analyzed by multiple, blinded raters and, where blinding was not possible (e.g. due to recognizable landmarks in the image stacks), all counts were validated by multiple raters. As an additional measure, only new sheaths with processes that could be traced back to the surviving oligodendrocyte cell body with Simple Neurite Tracer were included in our final dataset (Supplementary Fig. 3; Supplementary Video 3). Finally, myelin debris prevented faithful analysis of processes and sheaths from oligodendrocytes in the timepoint following cuprizone and therefore we excluded the timepoint immediately after the removal of cuprizone (i.e. 0d) from analysis.

#### Myelin sheath analysis of individual oligodendrocytes

*In vivo* z-stacks were collected from *MOBP-EGFP* mice using two-photon microscopy. Z-stacks were processed with a 1-pixel Gaussian blur filter to aid in the identification of myelin internodes. Myelin paranodes and nodes of Ranvier were identified as described previously^73^, by increase in fluorescence intensity for paranodes and a decrease to zero in EGFP fluorescence intensity for nodes. Myelin sheaths from surviving oligodendrocytes were traced using Simple Neurite Tracer at day −25 and day 21. In normal learning mice, two additional time points were traced that corresponded to day 0 and day 9 of training. To account for differences in measurement due to tracing, a subset of sheaths was traced five times at a single time point. Traces of the same sheath differed by less than 5.56 μm. Therefore, sheaths were defined as stable if their baseline and final lengths changed less than 5.56 μm. Sheaths that grew more than 5.56 μm were considered growing and those that shrank more than 5.56 μm were considered retracting. For myelin sheath analysis of surviving oligodendrocytes, surviving oligodendrocytes that resided within a volume of 425 × 425 × 100 μm^3^ from the pial surface were considered in Layer 1 and cells 100-336 μm^2^ were considered Layer 2/3. Myelin sheaths of surviving oligodendrocytes were tracked throughout time with the same FIJI scripts used for cell tracking. Only sheaths with visible processes back to the surviving cell body in at least one time point were counted. Sheaths were defined as new, lost, or persisting. Persisting sheaths lasted for the entire imaging time course; new sheaths appeared after day 0; and lost sheaths disappeared before the end of the imaging time course and were not visible for at least 2 consecutive timepoints. Our average total sheath count per surviving oligodendrocyte was 30.2±1.32. Assuming a normal range of 45±4 sheaths/oligodendrocyte^73^, our average sampling was 67-74% of all sheaths.

### Electrophysiology

Chronic *in vivo* recordings were conducted during 20 minute forelimb reach training sessions before, during, and after cuprizone treatment. A single 1.6mm vertical NeuroNexus recording electrode was chronically inserted into primary motor cortex (300μm anterior to bregma, 1.5mm lateral bregma) contralateral to the trained forelimb. Data was recorded using Cheetah acquisition software at 30kHz (NeuroNexus), and single unit activity was clustered using Spike Sort 3D (Neuralynx). Isolation Distance and L-Ratio was used to quantify cluster quality and noise contamination^81^. Spike data was binned at 10ms and trial-averaged. Heatmaps report average firing rate during 500 ms time window when the animal was not engaged in reach behavior.

### Statistics

A detailed and complete report of all statistics used, including definitions of individual measures, summary values, sample sizes, and all test results can be found in **Supplementary Table 2**. Sample sizes were not predetermined using statistical methods, but are comparable to relevant publications. All data were initially screened for outliers using IQR methods. Homozygous, C57BL/6N congenic *MOBP-EGFP* mice were bred with C57BL/6N mice to generate litters of experimental mice that were hemizygous for the *MOBP-EGFP* transgene. All mice in a litter underwent cranial window surgery, concurrent two-photon imaging timelines, and were designated to be a “batch”. When possible, experimental groups were replicated in multiple batches with multiple experimental groups per batch. However, due to the longitudinal nature of our study, and the clarity of cranial windows used to conduct two-photon imaging, we could not predict which mice would produce full datasets when assigning them to experimental conditions. As a result, not all batches contain all experimental groups, and we controlled for batch statistically. Statistical analyses were conducted using JMP (SAS) or MATLAB (MathWorks). We first assessed normality in all datasets using the Shapiro-Wilk test. When normality was violated, we used non-parametric rank-sum tests. When normality was satisfied, we used parametric statistics including paired or unpaired two-tailed Student’s t-tests (depending on within- or between-subjects nature of analysis), or one-way ANOVA with Tukey’s HSD post-hoc tests. Two-tailed tests and α ≤ 0.05 were always employed unless otherwise specified. For statistical mixed modelling, we used a restricted maximum likelihood (REML) approach with unbounded variance component, and least-squared Tukey’s HSD post-hoc tests. All models were conducted with either one or two fixed effects, in which case we ran full factorial models. For all models, we used “Mouse ID” as a random variable, and this random variable was nested within “batch” if data came from separate batches. Where we found significant effects, we subsequently calculated effect size using cohen’s d or hedge’s g, these results may be found in Supplementary Table 2. For data visualization, all error bars represent standard error on the mean unless otherwise specified. We used three-parameter (3P) logistic equation modelling to fit sigmoidal curves bound between an asymptote of 0 (baseline) and an estimated plateau to oligodendrocyte accumulation (either loss or gain) across time.

