## Supplementary Figures / Video legends for "Motor Learning Promotes Remyelination via New and Surviving Oligodendrocytes"

PRE-PRINT

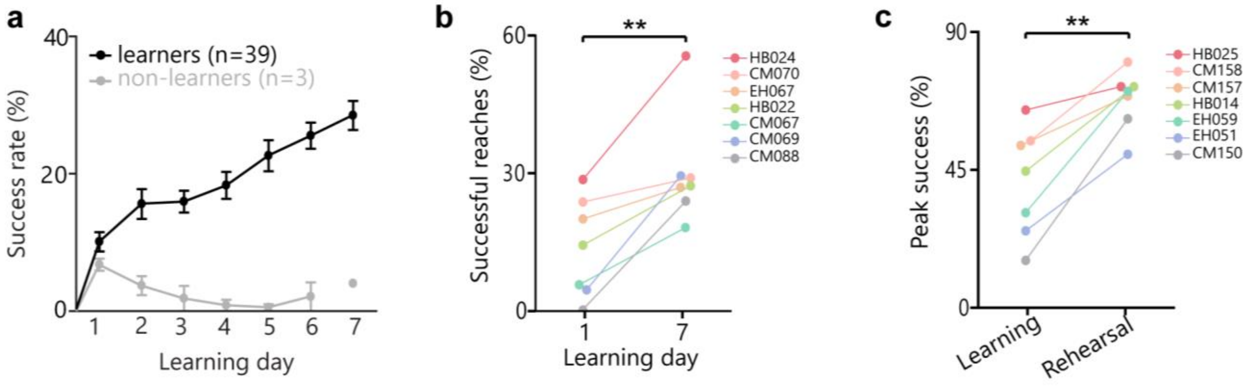


**Supplementary Fig. 1 | Learning and rehearsal of a forelimb reach task induce skill refinement. a,** A large majority (93%) of mice successfully learn to perform the forelimb reach task. “Learners” (black) gradually improve their reaching performance over the seven days of training, whereas “non-learners” (grey) show a progressive decrease in success rate and eventually stop making reach attempts around day 4. Note, the lone point in the “non-learner” group at day 7 is due to only one mouse making attempts on the last day of training. The other two mice had stopped trying. **b,** Successful reaches (%) significantly increase between learning days 1 and 7 (paired samples t-test; $t(6) = 4.80$, $p = 0.003$) for mice placed in “learning” group. **c,** Peak performance during rehearsal (successful reaches; %) is significantly higher than peak performance during learning (paired samples t-test; $t(6) = 5.47$, $p = 0.0016$) for mice placed in “rehearsal” group. Individual colors and traces reflect performance by individual mice.

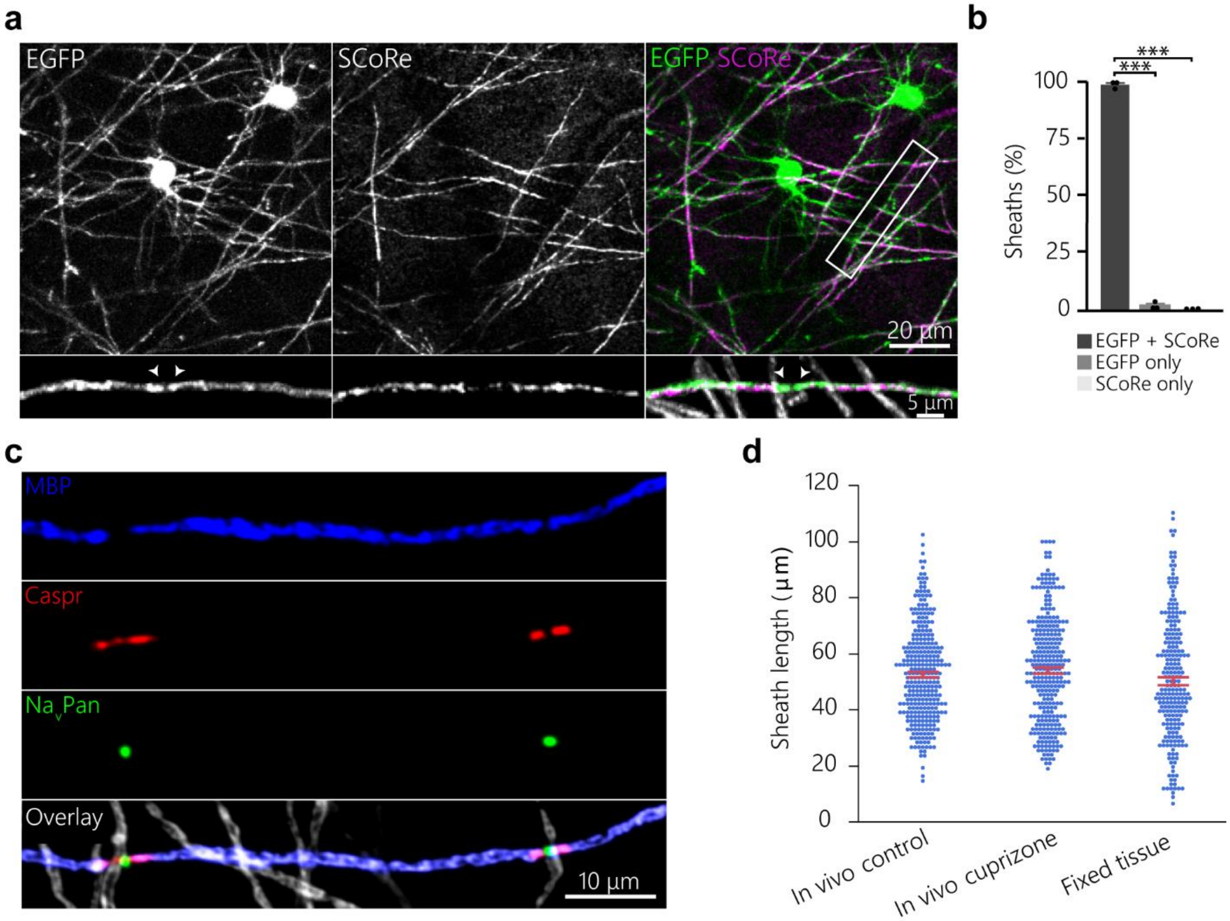


**Supplementary Fig. 2 | *In vivo* imaging of *MOBP-EGFP* faithfully reflects presence and length of myelin sheaths. a, b**, Maximum projections of cortical oligodendrocytes showing 98.24±0.92% colocalization of *in vivo MOBP-EGFP* and SCoRe signal in myelin sheaths, confirming *MOBP-EGFP* faithfully reflects presence of myelin (ANOVA, $F_{2,6}$=5596.220, p<0.0001). **c,** Maximum projection of 4% paraformaldehyde fixed tissue, stained for myelin (blue, MBP), paranodes (Caspr, red), and sodium channels (NavPan, green). **d,** No difference between sheath lengths measured using Simple Neurite Tracer in *in vivo* two-photon images of control and cuprizone-treated *MOBP-EGFP* mice, and in confocal images of sheaths immunostained for MBP in fixed tissue ($n_{sheaths}$=306, 297 and 233, respectively; points represent individual sheaths; ANOVA; $F_{2,833}$=2.53, p>0.08).

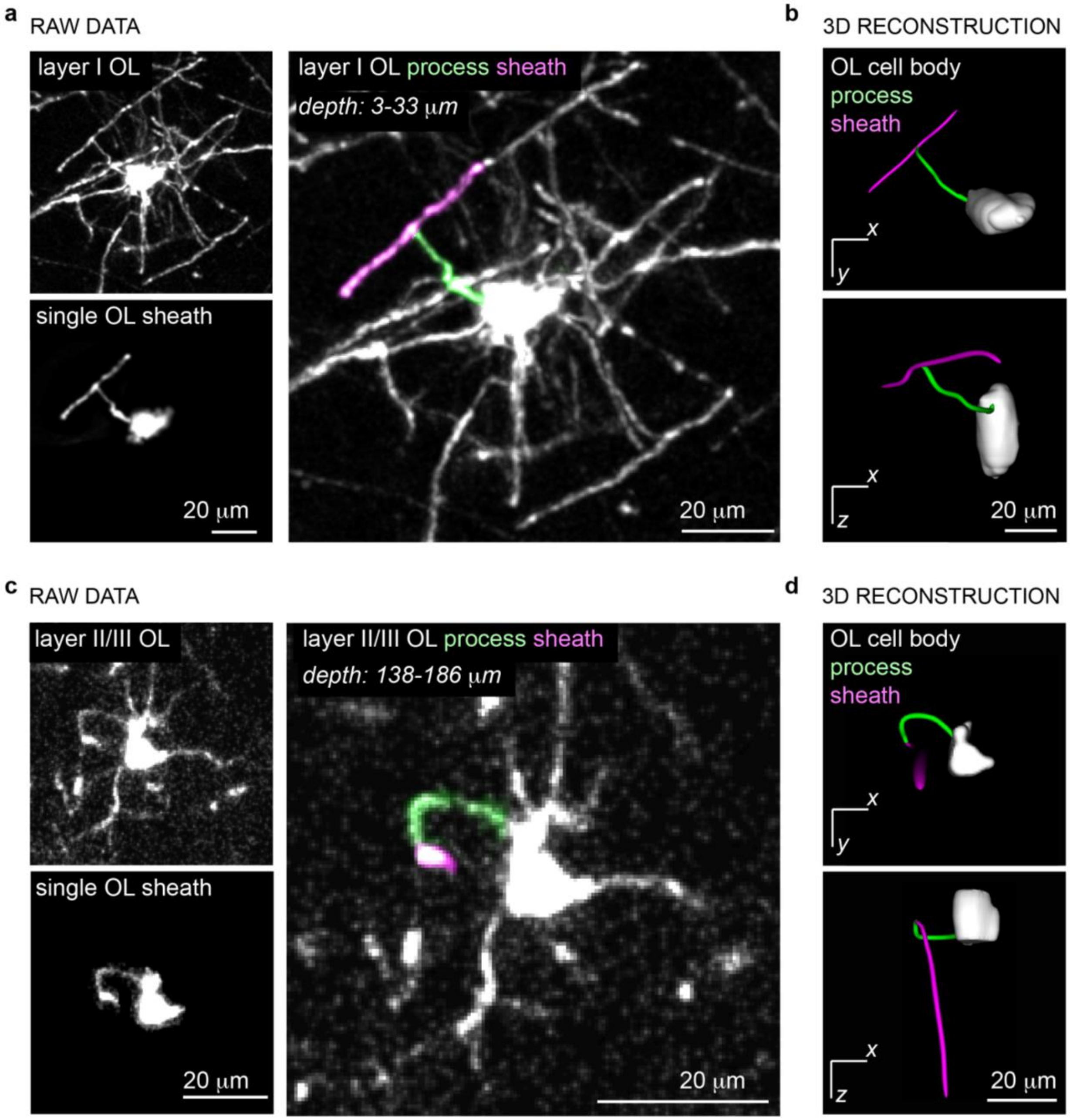


**Supplementary Fig. 3 | Semi-automated tracing with Simple Neurite Tracer faithfully reconstructs oligodendrocyte myelin sheaths and their connecting processes to the cell soma in layer I and layer II/III. a,** Maximum projection of a layer I oligodendrocyte (OL) imaged using *in vivo* two-photon microscopy, spanning a depth of 3-33 μm in motor cortex (top left), maximum projection of an isolated single sheath and process attached to the oligodendrocyte cell body (bottom left), and maximum projection and pseudo-colored sheath and process (right, sheath and process pseudo-colored). **b,** Three-dimensional (3D) reconstruction of the same layer I oligodendrocyte generated from the raw *in vivo* imaging data using the Simple Neurite Tracer plugin in FIJI. View of 3D volume in xy plane from below (top) and view of 3D volume through z (bottom). **c,** Maximum projection of a layer II/III oligodendrocyte imaged using *in vivo* two-photon microscopy, spanning a depth of 138-186 μm in motor cortex (top left), maximum projection of an isolated single sheath and process attached to the oligodendrocyte cell body (bottom left), and maximum projection and pseudo-colored sheath and process (right). **d,** 3D reconstruction of the same layer II/III oligodendrocyte generated from the raw *in vivo* imaging data using the Simple Neurite Tracer plugin in FIJI. View of 3D volume in xy plane (top) and view of 3D volume through z (bottom).

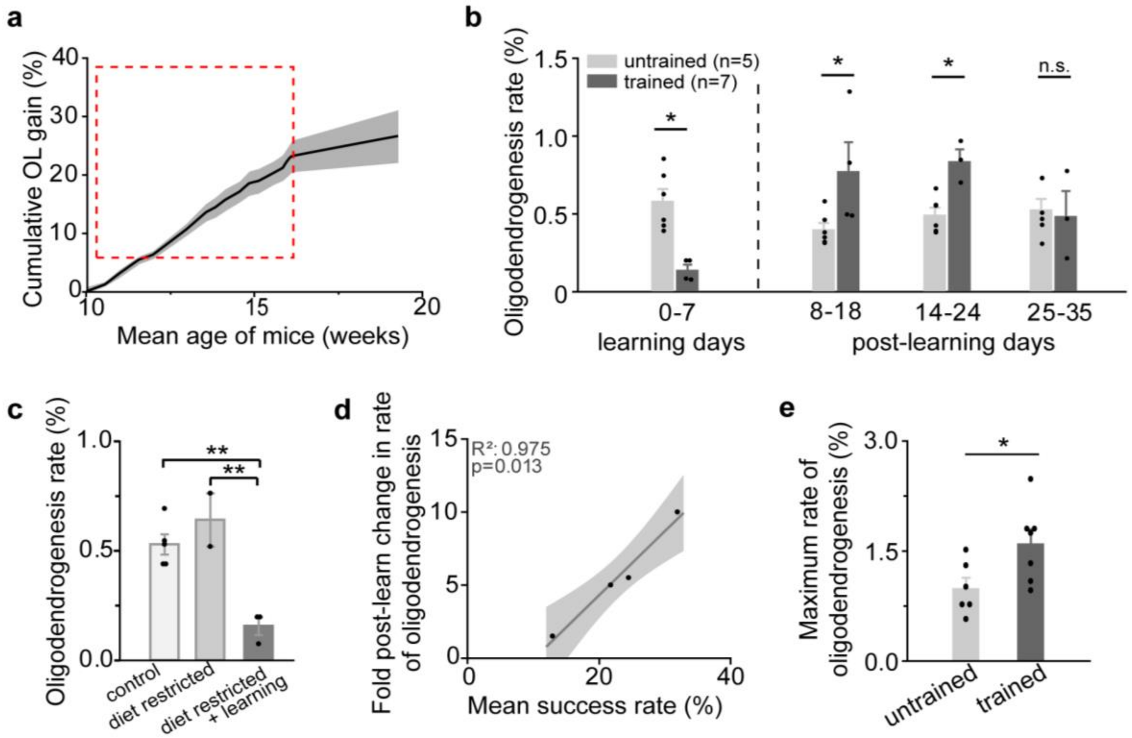


**Supplementary Fig. 4 | Dynamics of oligodendrocyte suppression and genesis following learning. a,** Overview of motor cortex oligodendrogenesis between the age of 10-20 weeks in n = 6 mice, showing a plateau in gain around 17 weeks. Red dashed box represents age during standard experimental timeline. Longitudinal time points are drawn beyond the red dashed box. **b,** Cross-sectional analyses show trained mice differ in rate of oligodendrogenesis during learning (Wilcoxon Rank-Sum, $p = 0.014$), day 8-18 post-learning ($p = 0.038$), and days 14-24 post-learning ($p = 0.024$). No differences are observed by days 25-35 post-learning ($p > 0.9$). Individual points represent individual mice. Trained group data from days 0-7 and 8-18 come from the standard "learning" group depicted in Fig 1b, and data from days 14-24 and 25-35 are drawn from the "rehearsal" group, following learning but before rehearsal. **c,** Main effect of diet restriction on rate of oligodendrogenesis (%; ANOVA; $F(2,8) = 18.13$, $p = 0.001$). Both diet-restricted and non-diet-restricted controls had higher rates of oligodendrocyte gain relative to diet-restricted mice that underwent learning (Tukey's HSD, $p = 0.001$ and $p = 0.005$, respectively). No difference in rate was found between fasting and no-fasting controls ($p > 0.1$). **d**, Mean success rate during learning is related to fold change in rate of oligodendrogenesis after learning (R-square = 0.98, $p = 0.01$). Line and shaded area represent line of fit and 95% confidence of fit. e, Trained mice (both from learning and rehearsal conditions) have increased maximum rates of oligodendrogenesis relative to controls ($t(10.61)=-2.49$, $p=0.03$).

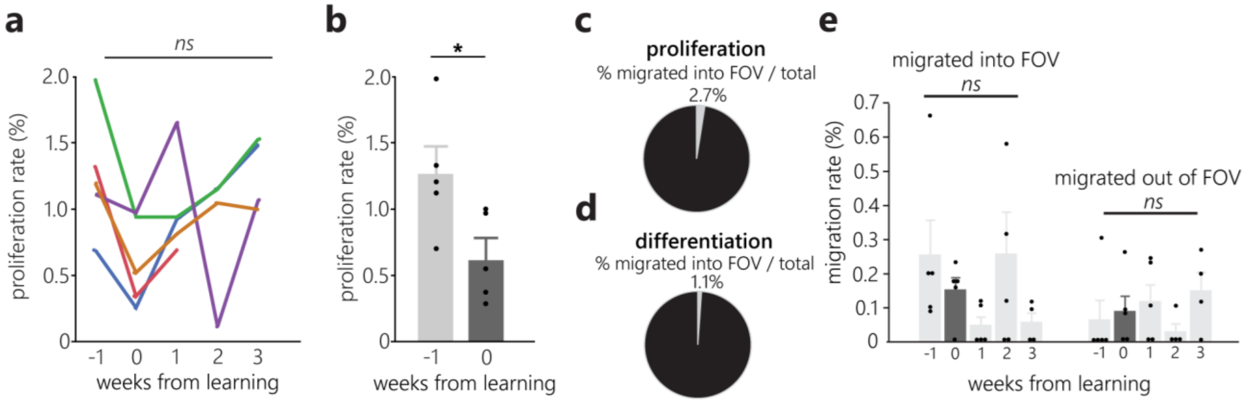


**Supplementary Fig. 5 | Effects of forelimb reach training on OPC proliferation and migration. a,** Proliferation data from Fig. 2d, where individual colors represent individual mice. All mice analyzed displayed a reduction in proliferation rate during learning, however, there was no main effect of time on proliferation rate - possibly due to high variability in rates following learning ($F(4,15)=2.341$). **b,** When compared cross-sectionally to the pre-learning phase, proliferation rate during learning (week 0) was decreased by more than half ($t(4) = -3.89$, $p = 0.018$; paired student's t-test). **c-d,** Only a small minority of proliferation and differentiation events occurred in OPCs that had migrated into the field of view throughout the course of the experiment (2.7% and 1.1%, respectively). **e,** No effect of learning on the rate of migration into or out of the field of view (FOV).

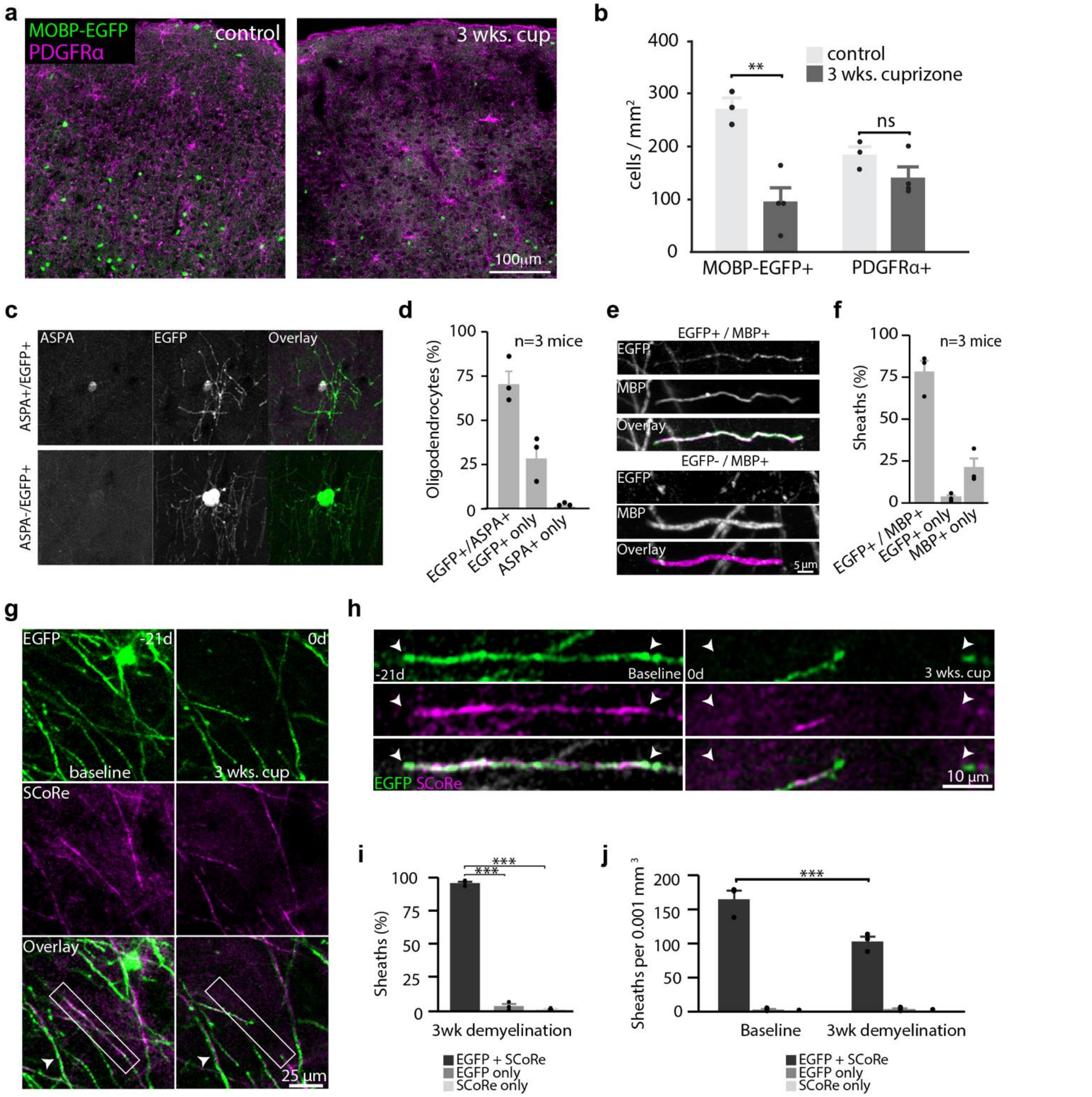


**Supplementary Fig. 6 | Cuprizone treatment results in loss of myelin and oligodendrocytes.**

**a, b,** Mean density of EGFP+ and PDGFRα+ cells in control and cuprizone-treated *MOBP-EGFP* mice; individual points represent individual mice. Interaction effect between drug (control vs. cuprizone) and cell type (EGFP+ vs PDGFRα+ ; ($F(1,5)=22.39$, $p=0.0052$) to predict cell density. While EGFP+ cell density is decreased in cuprizone-treated mice relative to controls (Tukey's HSD; $p=0.0086$), there is no difference in PDGFRα+ cells between control and cuprizone-demyelinated mice ($p>0.5$). **c,** Maximum projection of a $EGFP^+$, $ASPA^+$ oligodendrocyte (top) and a $EGFP^+$ and ASPA negative oligodendrocyte (bottom). Note the large size of the $ASPA^-/EGFP^+$ cell soma suggesting it is a recently born oligodendrocyte. **d,** After three weeks of cuprizone treatment, 70.73±12.78% of oligodendrocytes are both $EGFP^+$ and $ASPA^+$, 0.97±0.84% of cells are $ASPA^+$ and EGFP negative, while the remainder are $EGFP^+$ only ($n_{mice}$ = 3, $n_{cells}$ = 185). $EGFP^+$ only likely represent new oligodendrocytes that are in the early stages of the maturation process. **e,** Maximum projection of an $MBP^+$ myelin sheath with (top) and without EGFP (bottom) after three weeks of cuprizone. **f,** After three weeks of cuprizone, 83.27±1.76% of $MBP^+$ sheaths are also $EGFP^+$, and 14.85±0.79% of sheaths are $MBP^+$ and EGFP negative ($n_{mice}$ = 3, $n_{sheaths}$ = 351). **g,h** Maximum projections of cortical oligodendrocytes showing colocalization of myelin sheaths via *in vivo MOBP-EGFP* and SCoRe imaging both before cuprizone administration (-21 days) and immediately following its removal (0days). Note the surviving sheath (white arrow). **i,** Following 3 weeks of cuprizone diet, almost all myelin sheaths are visualized by both MOBP-EGFP and SCoRe (95.71±1.16; ANOVA, $F_{2,6}$=2012.94, p<0.0001). **j,** Cuprizone administration modulates sheath density ($F_{2,10}$=14.43, p=0.001). Cuprizone-fed mice have a reduced density of SCoRe+GFP positive sheaths relative to controls (p=0.0001), but no difference in GFP-only or SCoRe only sheaths.

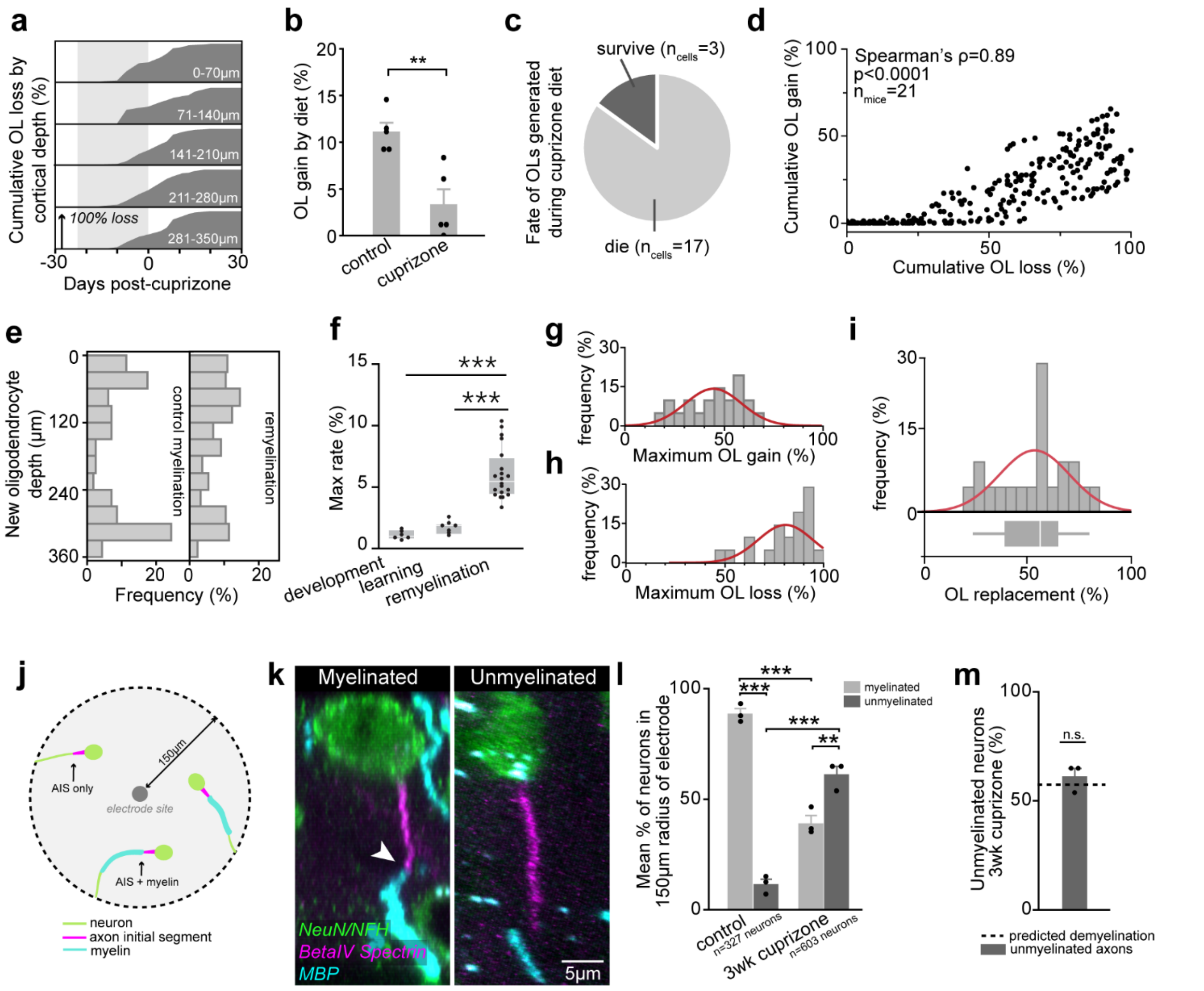


**Supplementary Fig. 7 | Dynamics of oligodendrogenesis and loss induced by cuprizone treatment. a,** Cumulative oligodendrocyte loss occurred evenly across cortical depths. Shaded area represents cuprizone diet. **b,** Oligodendrogenesis is suppressed during cuprizone diet relative to control diet ($t(6.54) = 4.10$, $p = 0.005$; Student's t-test). **c,** 85% of oligodendrocytes generated during cuprizone diet die within three weeks. **d,** Cumulative oligodendrocyte loss strongly predicts cumulative oligodendrocyte gain (Spearman's $\rho$=0.89). **e,** Oligodendrocytes generated during remyelination follow similar patterns of cortical depth to oligodendrocytes generated under normal conditions (Wilcoxon Rank-sum, $p > 0.1$). **f**, Maximum rates of oligodendrogenesis are modulated by remyelination ($F(29,2)=27.67$, $p<0.0001$; ANOVA). Rates are higher during remyelination than in healthy untrained and trained mice ($p<0.0001$ and $p<0.0001$, respectively; Tukey's HSD). **g, h, i,** Both maximum oligodendrocyte gain and loss vary strongly between mice, but normalizing gain to loss to obtain "oligodendrocyte replacement" (**i**) results in a normally distributed remyelination response. **j,k,** Representative diagram and images of peri-electrode immunohistochemistry analysis. Myelinated neurons within 150 microns of the electrode were co-labelled with MBP (myelin; cyan) beta-IV spectrin (axon initial segment (AIS); purple) and NeuN/NFH (neuron cell soma / distal axon; green; top), whereas unmyelinated axons did not co-localize with MBP (bottom). White arrowhead indicates a myelin sheath immediately proximal to an axon initial segment on a myelinated axon of a neuron in layer II/III in a XZ maximum projection. **l,** Cuprizone administration alters peri-probe axonal myelination (Two-way ANOVA; $F_{3,8}=110.51$, $p<0.0001$). Control mice have significantly more myelinated versus unmyelinated axons (Tukey's HSD, $p<0.0001$). At the cessation of cuprizone administration, cuprizone-fed mice have considerably fewer myelinated axons than controls ($p<0.0001$), and more unmyelinated than myelinated axons ($p=0.004$). As a result, cuprizone-fed mice have far more unmyelinated axons than healthy controls ($p<0.0001$). *Note: This shows loss of myelin proximal to the AIS, but myelin may be present elsewhere on the axon.* **m,** The proportion of unmyelinated neurons observed via IHC does not differ from the proportion of myelin loss predicted by sigmoidal demyelination characterized in **Fig. 3e-h** (one-sample t-test, $t(2)=1.10$, $p>0.3$).

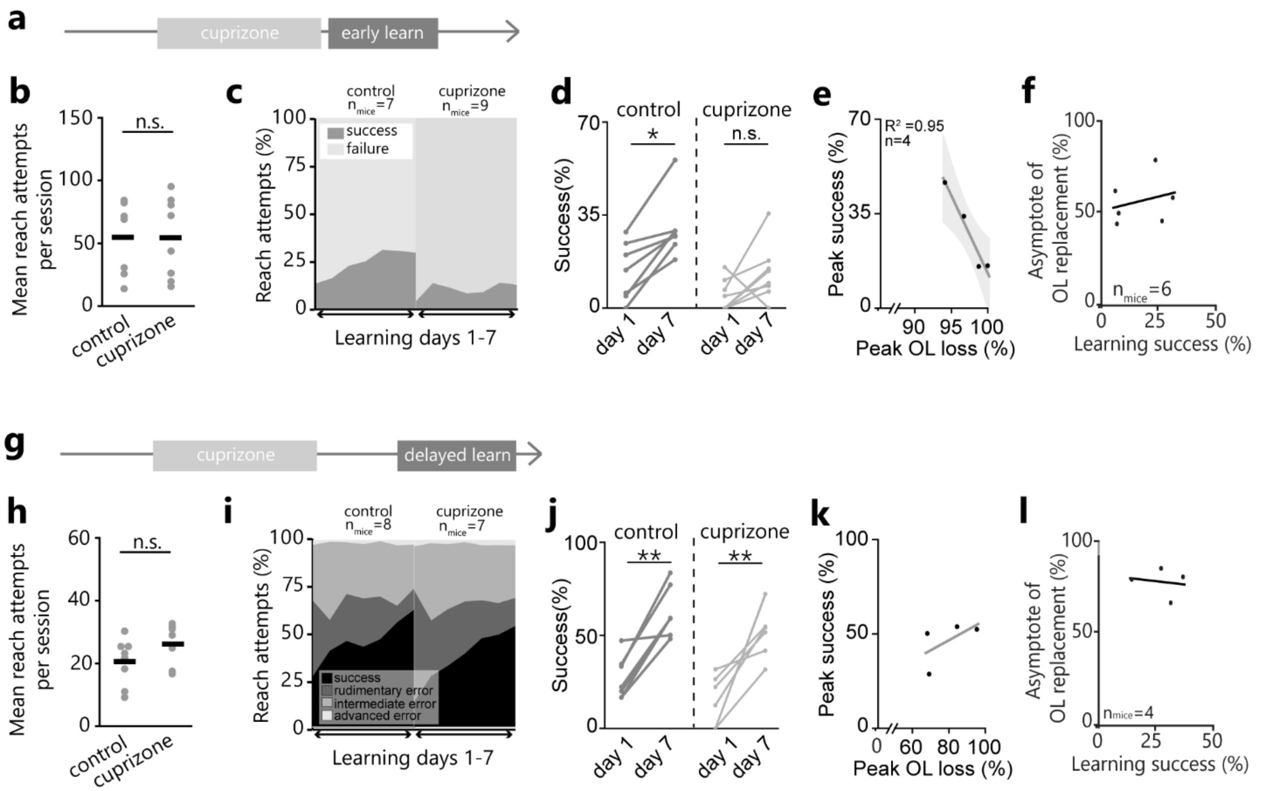


**Supplementary Fig. 8 | Demyelination induces deficits in early motor learning**. **a,** Timeline for learning forelimb reach task after cuprizone. **b**, No difference in mean reach attempts per session during early-learning between control and cuprizone-treated mice (Student's t-test, t(12.95) = 0.05, p > 0.9). **c,** Area plot of reach attempt outcome (success vs. failure) across forelimb reach learning days in both control and cuprizone-treated mice. **d,** Control mice have improved success rates day 7 of training relative to day 1 (Paired Student's T-test; t(6)=4.7, p=0.003), but cuprizone-treated mice do not (t(7)=1.96). **e,** Maximum oligodendrocyte loss is related to peak performance during training ($R^2$=0.95, p=0.02; line and shaded area represent line of fit and 95% confidence of fit). **f,** No relationship between mean learning success rate (%) and asymptote of oligodendrocyte replacement in early learners. **h,** No difference in mean reach attempts per session during delayed-learning between control and cuprizone-treated mice (Student's t-test, t(12.95)=1.54, p>0.1). **j**, Both control and cuprizone-treated mice improve their reaching success between days 1 and 7 of delayed-learning (Paired student's t-test, p=0.0005 and p=0.004, respectively). **k**, No relationship between maximum oligodendrocyte loss and reaching performance during delayed learning. **l,** No relationship between delayed learning success rate and asymptote of oligodendrocyte replacement post-cuprizone.

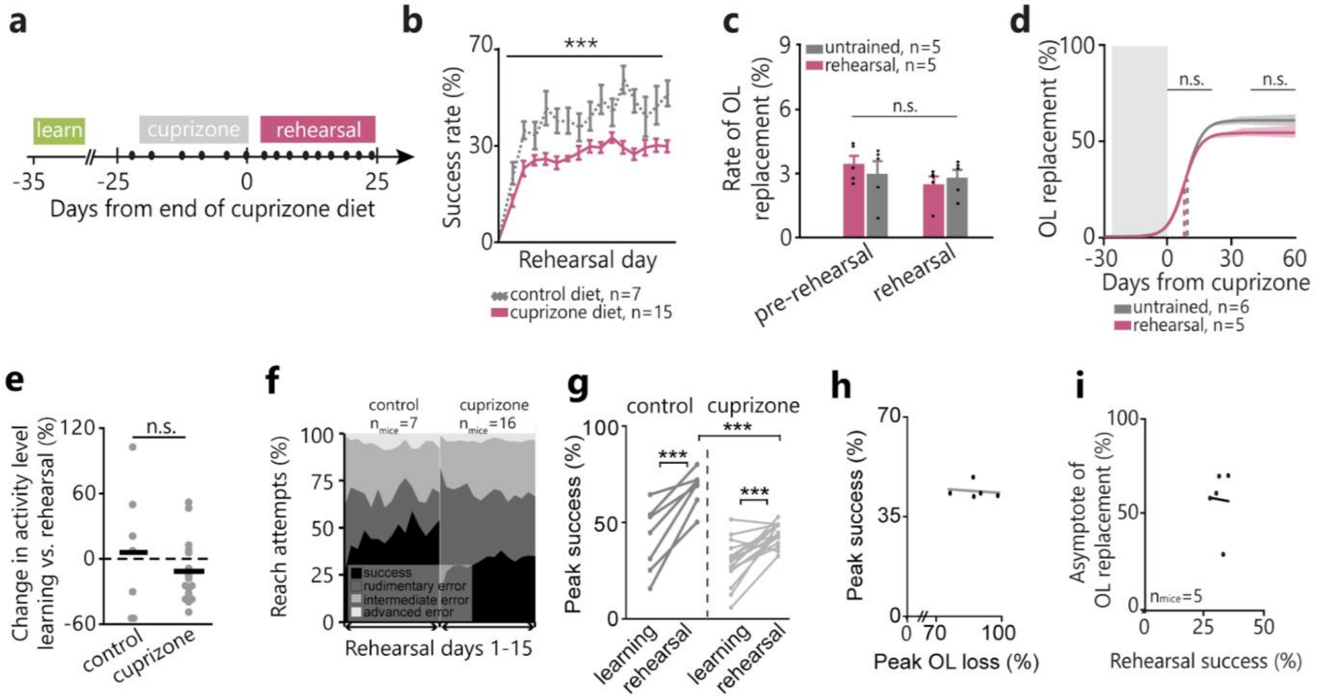


**Supplementary Fig. 9 | Motor skill rehearsal does not modulate remyelination. a,** Timeline of reach task rehearsal post cuprizone diet. **b**, Main effect of drug on reaching success during rehearsal (F(1,14)=27.73, p<0.0001). **c,d**, No effect of rehearsal on rate, inflection point, or asymptote of oligodendrocyte replacement. **e,** No effect of cuprizone on change in reaching behavior between learning and rehearsal. **g,** Interaction effect between performance phase (learning vs. rehearsal) and drug (control vs. cuprizone) to predict success rate (F(1)=4.62, p=0.04). While control and cuprizone mice do not differ in success rate during pre-cuprizone learning, control mice perform significantly better during rehearsal relative to cuprizone-treated mice (Tukey's HSD, p=0.0004). Both cuprizone and cuprizone-treated mice have improved performance during rehearsal relative to learning (p=0.0001 and p<0.0001, respectively). **h**, No relationship between peak oligodendrocyte loss post-cuprizone and peak reaching success rate during rehearsal.

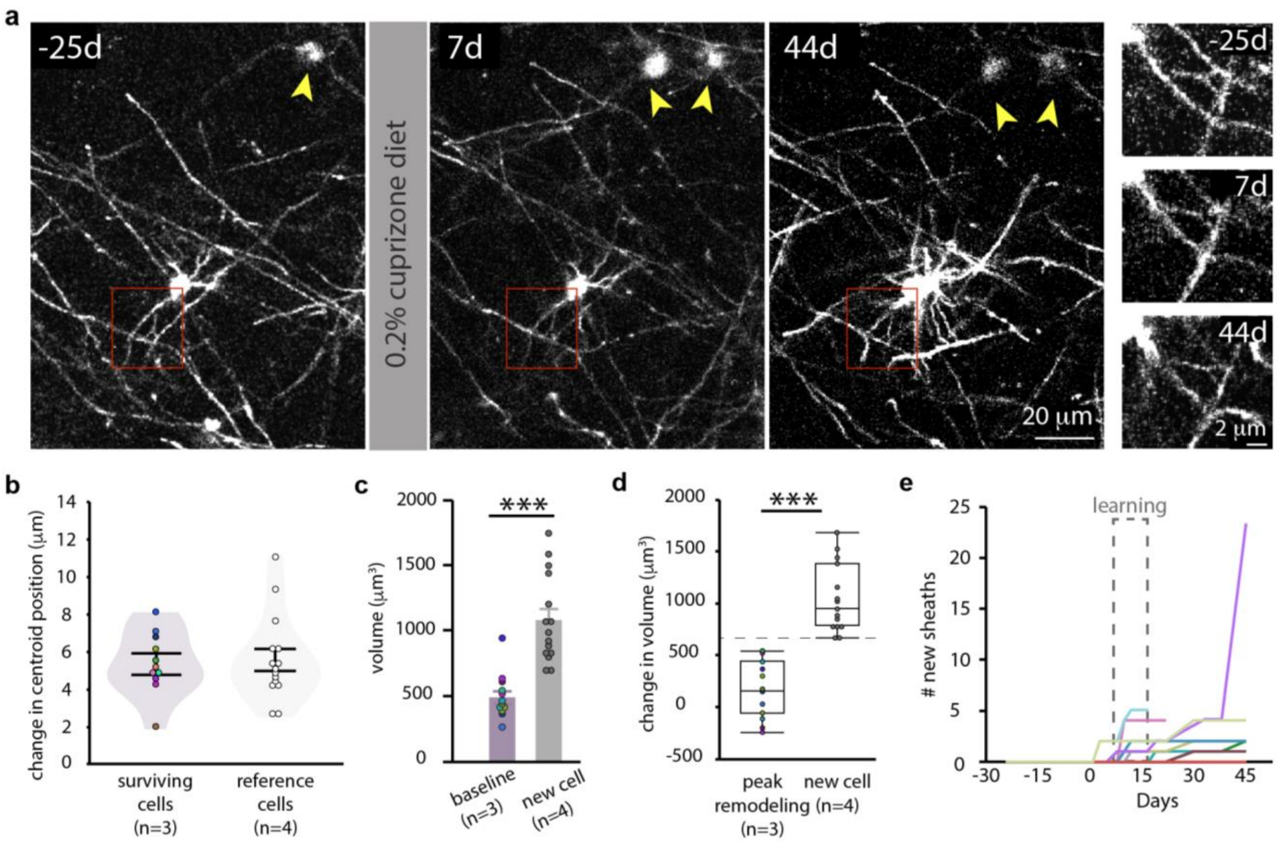


**Supplementary Fig. 10 | Identification of oligodendrocytes that survive demyelination. a,** Representative image outlining the methodology for following surviving oligodendrocytes over time. Single plane image of the same oligodendrocyte at baseline (-25d), one week after demyelination (7d), and six weeks after demyelination (44d). Red boxes highlight one example of the same oligodendrocyte processes lasting for the duration of the study. The maintenance of the spatial relationship between the oligodendrocyte of interest and other oligodendrocytes in the field of view (yellow arrowheads) provide further confirmation of oligodendrocyte identity. Note the new cell that appears at 7d. **b,** Change in centroid position of reference oligodendrocytes within the z-stack and surviving cell bodies from baseline to day of peak remodeling—i.e. the day where the largest number of sheaths were added by a given oligodendrocyte. **c,** Surviving oligodendrocytes at baseline are significantly smaller than new oligodendrocytes ($t(21.91) = -5.81$, $p < 0.0001$, Student's t-test). **d,** Change in volume of surviving oligodendrocytes from baseline to peak remodeling is significantly smaller than the volume of new oligodendrocytes ($t(23.88 = -7.59$, $p < 0.0001$). **e,** Dynamics of sheath addition over time. Each line represents an individual oligodendrocyte.

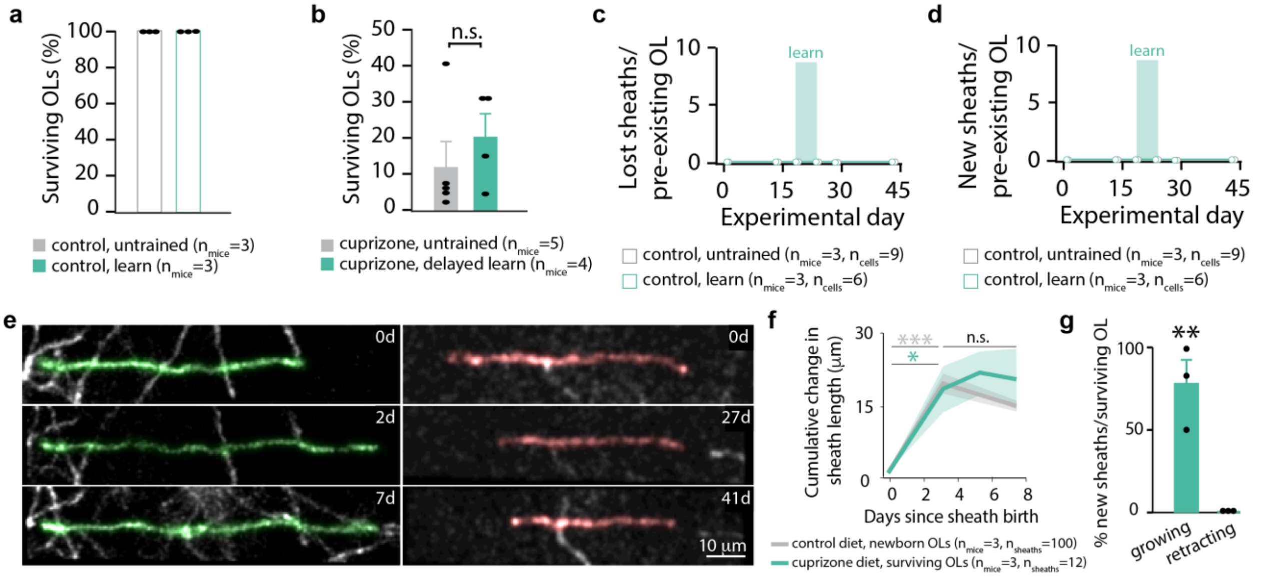


**Supplementary Fig. 11 | Dynamics of myelin sheath generation by pre-existing oligodendrocytes. a,** No oligodendrocytes are lost in healthy mice. **b,** No difference in percent of oligodendrocytes (OLs) surviving demyelination in untrained and delayed learning groups (Wilcoxon Rank-Sum, $p>0.5$). **c, d,** No sheaths are lost (c) nor generated (d) on mature oligodendrocytes in healthy trained or untrained conditions. **e,** Maximum projection of new sheaths generated after cuprizone exhibiting growth (pseudocolored green, left) and retraction (pseudocolored red, right). **f,** New myelin sheaths change in length in the week following their generation, whether they are from new oligodendrocytes (control: $F(3,302)=47.94$, $p<0.0001$) or from surviving oligodendrocytes after cuprizone-demyelination (cuprizone diet: $F(3,29)=5.31$, $p=0.0049$). Sheaths in both control and cuprizone treatment stabilize their length within 3 days of sheath birth (d0 vs. d3, $p<0.0001$ in control and $p=0.028$ in cuprizone; Tukey's HSD). Line and shading represent mean and SEM. **g,** Sheaths from pre-existing oligodendrocytes grow more often than they retract the first three days post-generation (Wilcoxon Rank-Sum, $p=0.0029$).

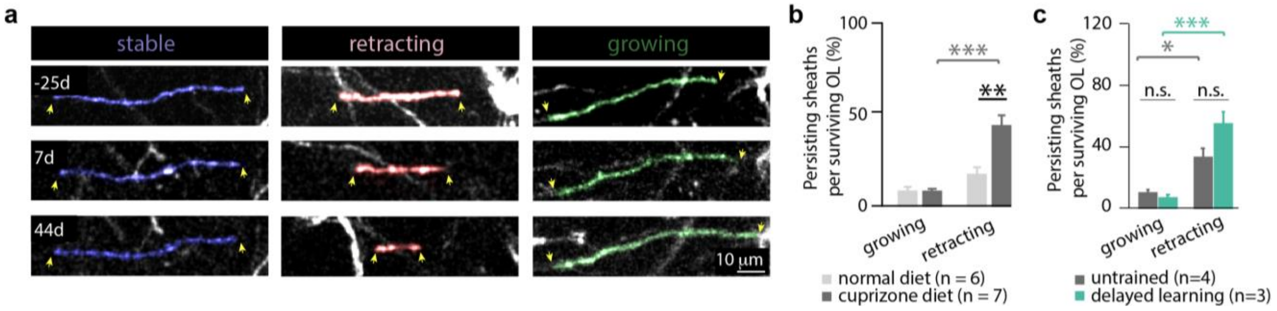


**Supplementary Fig. 12 | Surviving oligodendrocyte pre-existing myelin sheath dynamics during remyelination. a**, Behavior of pre-existing myelin sheaths that persist throughout study. Relevant sheaths are pseudocolored. **b**, Three weeks into remyelination, sheath retraction is significantly increased ($F(3,22) = 18.65$, $p < 0.0001$) when compared to age-matched controls (Tukey's HSD, $p = 0.0006$) and when compared to the percent of sheaths growing in cuprizone-treated mice ($p < 0.0001$). **c,** No effect of delayed learning on sheath dynamics during remyelination. Sheaths retract more than they grow in both untrained ($p = 0.016$) and delayed learning mice ($p = 0.0003$).

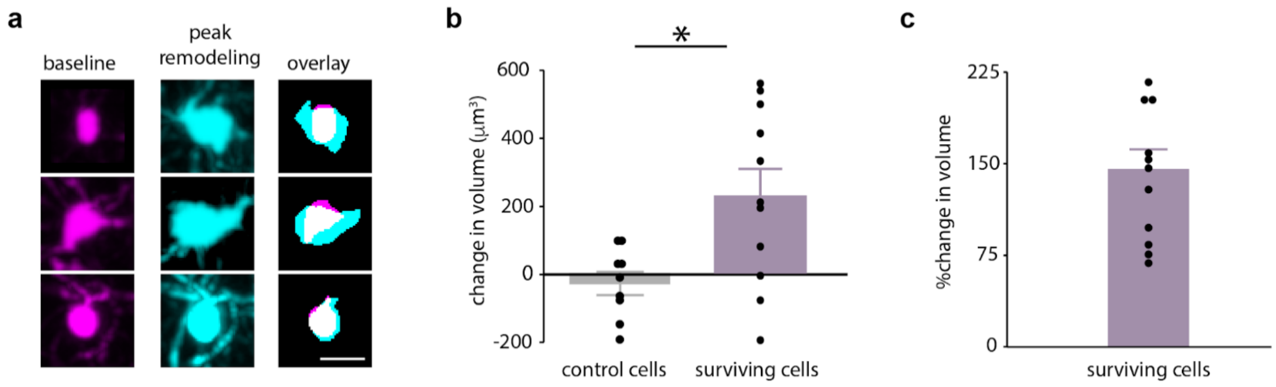


**Supplementary Fig. 13 | Surviving oligodendrocyte cell soma volume changes during remyelination. a,** Maximum projection of surviving oligodendrocyte cell bodies at baseline (left, magenta), peak remodeling (middle, cyan), and overlaid (right). Scale bar is 10 µm. **b,** Oligodendrocytes in normal untrained mice display little change in cell body volume throughout the study, from baseline (0d) to 43d. Surviving cells in delayed learning mice show dramatic increase in cell soma volume from baseline to day of peak remodeling when compared to oligodendrocytes in normal untrained mice ($t(12.24) = 2.56$, $p = 0.025$, Student's t-test). **c,** Percent change in volume between baseline and day of sheath addition for surviving cells engaging in remodeling.

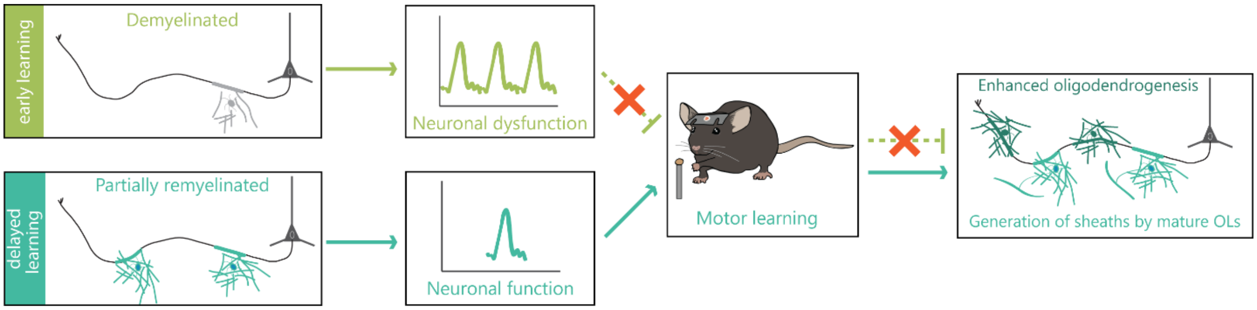


**Supplementary Fig. 14 | Summary diagram of effects of motor learning on oligodendrogenesis during remyelination.** In green, demyelinated mice are shown to have neuronal dysfunction (ie, hyperexcitability), and are unable to learn the forelimb reach task. They therefore receive no benefit of behavioral intervention during remyelination. In blue, partially remyelinated mice have neuronal function that is indistinguishable from healthy controls. When trained at this partial remyelination timepoint, mice are able to learn the forelimb reach task which enhances oligodendrogenesis and promotes the generation of new sheaths by surviving oligodendrocytes.

### SUPPLEMENTARY VIDEO LEGENDS

**Supplementary Video 1 | Categorization of reach attempts during forelimb reach training.** Categorization of four different possible outcomes of reach attempts during forelimb reach task. Success involves correct targeting, grasping, and retrieval of the pellet inside the training box. Reach error (rudimentary) involves incorrect targeting of the pellet – no contact with the pellet is made. During a grasp error (intermediate), the mouse correctly reaches for the pellet but does not successfully grasp its hand around the pellet. During a retrieval error (advanced), the mouse is able to reach and grasp onto the pellet, but does not successfully retrieve it into the box.

**Supplementary Video 2 | Developmental oligodendrogenesis in motor cortex.** Maximum intensity projection of 0-336µm of forelimb region of motor cortex across 43 days of two-photon *in vivo* imaging in a *MOBP-EGFP* mouse implanted with a cranial window. New EGFP-positive oligodendrocytes (labelled in green) are generated across scope of experiment, when mouse is aged 10-17 weeks. Developmental oligodendrogenesis begins to slow after 16 weeks of age. (Frame rate: 2 frames per second).

**Supplementary Video 3 | Using Simple Neurite Tracer to trace the majority of the oligodendrocyte arbor.** *In vivo* two-photon imaging of sheaths from a mature oligodendrocyte. This field was acquired in motor cortex of an *MOBP-EGFP* mouse implanted with a cranial window (P63; depth = 3-51 µm) and follows all traceable processes through z in both single plane images and with reconstructions. Note yellow arrow pointing to node of Ranvier and rotating 3D reconstruction of oligodendrocyte cell body (white), sheaths (magenta), and processes (green). (Frame rate: 15 frames per second).

**Supplementary Video 4 | Categorization of existing sheath dynamics: retracting.** *In vivo* two-photon imaging of the retraction of a myelin sheath from an existing oligodendrocyte in the motor cortex of an *MOBP-EGFP* mouse implanted with a cranial window (P69; depth=3–9 µm). The sheath begins continuous retraction within the first week (7d). Note the extended time period for which the sheath is retracting (7d-39d). (Frame rate: 2 frames per second).

**Supplementary Video 5 | Categorization of existing sheath dynamics: growing.** *In vivo* two-photon imaging of the growth of a myelin sheath from an existing oligodendrocyte in the motor cortex of an *MOBP-EGFP* mouse implanted with a cranial window (P69; depth = 9–18 µm). This myelin sheath (green) is stable for 43 days, then extends over the next 19 days. (Frame rate: 2 frames per second).

**Supplementary Video 6 | Dynamics of demyelination and remyelination.** *In vivo* two-photon imaging of a field undergoing de- and remyelination in motor cortex of a *MOBP-EGFP* mouse implanted with a cranial window (P68; depth = 0–336 µm) and treated with 0.2% cuprizone diet for three weeks. Images were acquired every 2–3 days for 43 days. In this 3-week partial demyelination model, ~90% of EGFP-positive expressing oligodendrocytes are lost relative to baseline (-21d) number and ~50% of oligodendrocytes are regained three weeks into remyelination (21d). (Frame rate: 2 frames per second).

**Supplementary Video 7 | Cuprizone-mediated myelin and oligodendrocyte loss.** *In vivo* two-photon imaging of an individual oligodendrocyte undergoing demyelination and death in motor cortex of a *MOBP-EGFP* mouse implanted with a cranial window (P63; depth = 3–18 µm) and treated with 0.2% cuprizone diet for three weeks. Images were acquired every 2–3 days for 43 days. Note loss of EGFP+ myelin sheaths at the end of cuprizone treatment (0d) and loss of cell body 1.5 weeks later (11d). (Frame rate: 2 frames per second).

**Supplementary Video 8 | Myelin sheath generation by a new oligodendrocyte in the healthy brain.** *In vivo* two-photon imaging of the myelin sheaths from a newly generated oligodendrocyte in a healthy mouse, as well as semi-automated traces which highlight the connection to the oligodendrocyte cell body for each sheath counted. This field was acquired in the motor cortex of an *MOBP-EGFP* mouse implanted with a cranial window (P67; depth = 3-57 µm). This video contains two time-points: two days pre-generation (-2d), 7 days post-generation (7d). This newly generated oligodendrocyte forms a total of 40 new sheaths, 36 days after baseline imaging. This video is divided into three parts: first, maximum projections of each time point of the entire 54 µm z-stack, which correspond to the figure images in Fig. 4a (top, control diet); second, single plane images of the z dimension; and third, the same images overlaid with semi-automated tracings. These semi-automated tracings were used to verify the connection between newborn sheaths and the newborn oligodendrocyte cell body. (Frame rate: 2 frames per second).

**Supplementary Video 9 | Increased myelin sheath generation by a newly generated oligodendrocyte during remyelination.** *In vivo* two-photon imaging of the myelin sheaths from a newly generated oligodendrocyte during the first week of remyelination, as well as semi-automated traces which highlight the connection to the oligodendrocyte cell body for each sheath counted. This field was acquired in the motor cortex of an *MOBP-EGFP* mouse implanted with a cranial window (P64; depth = 21-111 μm). This video contains two time-points: two days pre-generation (-2d), 7 days post-generation (7d). This oligodendrocyte is generated during the first week of remyelination, 4 days after the cessation of cuprizone diet, and forms a total of 53 new sheaths. This video is divided into three parts: first, maximum projections of each time point of the entire 90 μm z-stack which correspond to the figure images in Fig. 4a (bottom, cuprizone diet); second, single plane images of the z dimension; and third, the same images overlaid with semi-automated tracings. These semi-automated tracings verify the connection between new sheaths and the oligodendrocyte cell body. (Frame rate: 2 frames per second).

**Supplementary Video 10 | Generation of a remodeling myelin sheath by a newly generated oligodendrocyte during remyelination.** *In vivo* two-photon imaging of the generation of a remodeling myelin sheath from a newly generated oligodendrocyte during the second week of remyelination, as well as semi-automated traces of the connection to the oligodendrocyte cell body for this myelin sheath. This field was acquired in the motor cortex of an *MOBP-EGFP* mouse implanted with a cranial window (P65; depth = 15-24 μm). This video contains three time points: baseline (-25d; note unmyelinated section), 4 days post-cuprizone (4d), and 11 days post-cuprizone (11d; note new oligodendrocyte and sheath). This oligodendrocyte was generated during the second week of remyelination and forms a new sheath on a never-before myelinated location. Yellow arrow corresponds to the eventual location of the junction between the new oligodendrocyte process and the new remodeling sheath. This video is divided into three parts: first, maximum projections of each time point of the entire 9 μm z-stack, which corresponds to Fig. 4h (top, remodeling); second, single plane images of the z dimension, with relevant sheaths pseudo-colored; and third, the same images overlaid with semi-automated tracings. These semi-automated tracings verify the connection between the new remodeling sheath and the new oligodendrocyte cell body. (Frame rate: 2 frames per second).

**Supplementary Video 11 | Generation of a remyelinating sheath by a new oligodendrocyte during remyelination.** *In vivo* two-photon imaging of the generation of a remyelinating myelin sheath from a new oligodendrocyte during the second week of remyelination, as well as semi-automated traces of the connection to the oligodendrocyte cell body for relevant sheaths. This field was acquired in the motor cortex of an *MOBP-EGFP* mouse implanted with a cranial window (P65; depth = 0-24 μm). This video contains three time points: baseline (-25d; note original oligodendrocyte and sheath, pseudocolored pink), 7 days post-cuprizone (7d; note loss of baseline oligodendrocyte), and 11 days post-cuprizone (11d; note new oligodendrocyte and sheath, pseudocolored yellow). The original myelin sheath and oligodendrocyte are lost in between 4 and 7 days after the cessation of cuprizone, and this location is then immediately remyelinated by a new oligodendrocyte, generated 7-9 days after the cessation of cuprizone diet. This video is divided into three parts: first, maximum projections of each time point of the entire 24 μm z-stack, which correspond to the figure images in Fig. 4h (bottom, remyelinating); second, single plane images of the z dimension, with relevant sheaths pseudo-colored; and third, the same images overlaid with semi-automated tracings through z. These semi-automated tracings verify the connection between the original oligodendrocyte cell body and original myelin sheath, as well as the connection between the new remyelinating sheath and the oligodendrocyte cell body. (Frame rate: 2 frames per second).

**Supplementary Video 12 | Loss of a pre-existing myelin sheath on a surviving oligodendrocyte.** *In vivo* two-photon imaging of the loss of a pre-existing myelin sheath from a surviving oligodendrocyte during and after the administration of cuprizone diet, as well as semi-automated traces of the connection to the oligodendrocyte cell body for this sheath. This field was acquired in the motor cortex of an *MOBP-EGFP* mouse implanted with a cranial window (P63; depth = 45-54 μm). This video contains three time points: baseline (-25d), 11 days pre-cuprizone cessation (-11d), and 44 days post-cuprizone (44d). Myelin sheath loss is a protracted process that in this example occurred over 29 days. This video is divided into three parts: first, maximum projections of each time point of the entire 9 μm z-stack, which correspond to the figure images in Fig. 7b; second, single plane images of the z dimension, with relevant sheaths pseudo-colored; and third, the same images overlaid with semi-automated tracings. These semi-automated tracings verify the connection between the surviving oligodendrocyte cell body and the lost myelin sheath. (Frame rate: 2 frames per second).

**Supplementary Video 13 | Generation of a new myelin sheath by a surviving oligodendrocyte.** *In vivo* two-photon imaging of the generation of a new myelin sheath from a surviving oligodendrocyte during remyelination, as well as semi-automated traces of the connection to the oligodendrocyte cell body for this sheath. This field was acquired in the motor cortex of an *MOBP-EGFP* mouse implanted with a cranial window (P63; depth = 51-66 μm). In contrast to sheath loss, sheath addition is a rapid process that results in a stable sheath within 3 days of initial myelin generation. This video contains four time-points: baseline (-25d), 11 days post-cuprizone (11d; note extensive sheath loss and presence of myelin debris), 18 days post-cuprizone (18d; note brightening of oligodendrocyte cell body and cessation of sheath loss), and 44 days post-cuprizone (44d; note addition of new sheaths by the surviving oligodendrocyte). This video is divided into three parts: first, maximum projections of each time point of the entire 15 μm z-stack, which correspond to the figure images in Fig. 7c; second, single plane images of the z dimension, with relevant sheaths pseudo-colored; and third, the same images overlaid with semi-automated tracings. These semi-automated tracings verify the connection between the surviving oligodendrocyte cell body and the new myelin sheath. (Frame rate: 2 frames per second).

**Supplementary Video 14 | Extended generation of new sheaths by a surviving oligodendrocyte.** *In vivo* two-photon imaging of the generation of myelin sheaths from a pre-existing oligodendrocyte during remyelination. This field was acquired in the motor cortex of an *MOBP-EGFP* mouse implanted with a cranial window (P63; depth = 3-42 μm). This video contains eight time points: baseline (-25d), 14 days pre-cuprizone cessation (-14d), 4 days pre-cuprizone cessation (-4d), 11 days post-cuprizone (11d), 18 days post-cuprizone (18d), 28 days post-cuprizone (28d), 35 days post-cuprizone (35d), and 44 days post-cuprizone (44d). This oligodendrocyte began generating new myelin sheaths (green) on day 3 of learning (11d) and continues over 33 additional days (44dNote that this surviving oligodendrocyte retains a number of sheaths (magenta, pre-existing) for the duration of the imaging time period, in addition to losing sheaths and gaining new sheaths (green, new). This video is divided into two parts. The first part shows single plane images of the z dimension, overlaid with semi-automated tracings. These semi-automated tracings verify the connection between the surviving oligodendrocyte cell body, its processes, and their associated myelin sheaths. The second part shows 3D reconstructions of these semi-automated traces over time. In this section, we show reconstructions of the processes and sheaths with verified connections to the surviving oligodendrocyte cell body over 69 days. (Frame rate: 2 frames per second). *N.B. Cell body from 44 days post-cuprizone used in all time points for reconstruction consistency.*

**Supplementary Video 15 | Generation of a new myelin sheath by a surviving oligodendrocyte in Layer 2/3.** *In vivo* two-photon imaging of the generation of a new myelin sheath from a surviving oligodendrocyte in Layer 2/3 during remyelination, as well as semi-automated traces of the connection to the oligodendrocyte cell body for this sheath. This field was acquired in the motor cortex of an *MOBP-EGFP* mouse implanted with a cranial window (P63; depth = 183-198 μm). This video contains three time-points: baseline (-25d), 5 days post-cuprizone (5d), and 7 days post-cuprizone (7d; note new process and myelin sheath). Note that this sheath runs perpendicular to the imaging plane, in contrast to sheaths generated by Layer 1 oligodendrocytes, which run parallel to the imaging plane. This video is divided into three parts: first, maximum projections of each time point of the entire 15 μm z-stack, which correspond to the figure images in Fig. 7k; second, single plane images of the z dimension, with relevant sheaths pseudo-colored; and third, the same images overlaid with semi-automated tracings. These semi-automated tracings verify the connection between the Layer 2/3 surviving oligodendrocyte cell body and the newly generated myelin sheath. (Frame rate: 2 frames per second).

**Supplementary Video 16 | Generation of a new remodeling sheath by a surviving oligodendrocyte.** *In vivo* two-photon imaging of the generation of a new, remodeling myelin sheath from a surviving oligodendrocyte during remyelination, as well as semi-automated traces of the connection to the oligodendrocyte cell body for this sheath. This field was acquired in the motor cortex of an *MOBP-EGFP* mouse implanted with a cranial window (P63; depth = 6-18 μm). This video contains three time-points: baseline (-25d), 28 days post-cuprizone (28d; note sheath loss), and 44 days post-cuprizone (44d; note sheath generation). This video is divided into three parts: first, maximum projections of each time point of the entire 12 μm z-stack, which correspond to the figure images in Fig. 7m (top, remodeling); second, single plane images of the z dimension, with relevant sheaths pseudo-colored; and third, the same images overlaid with semi-automated tracings. These semi-automated tracings verify the connection between the surviving oligodendrocyte cell body and the newly generated, remodeling myelin sheath. (Frame rate: 2 frames per second).

**Supplementary Video 17 | Generation of a new remyelinating sheath by a surviving oligodendrocyte.** *In vivo* two-photon imaging of sheath loss and subsequent remyelination by the same surviving oligodendrocyte, as well as semi-automated traces of the connection to the oligodendrocyte cell body for relevant sheaths. This video contains three time-points: baseline (-25d), 28 days post-cuprizone (28d; note sheath loss), and 44 days post-cuprizone (44d; note sheath replacement with new connecting process). Sheath loss occurred over a period of two weeks following the cessation of the cuprizone diet and, once the sheath was lost by the surviving oligodendrocyte, this same surviving oligodendrocyte replaced the sheath within a week. This field was acquired in the motor cortex of an *MOBP-EGFP* mouse implanted with a cranial window (P63; depth = 12-18 μm). This video is divided into three parts: first, maximum projections of each time point of the entire 12 μm z-stack, which correspond to the figure images in Fig. 7m (bottom, remyelinating); second, single plane images of the z dimension, with relevant sheaths pseudo-colored; and third, the same images overlaid with semi-automated tracings. These semi-automated tracings verify the connection between the surviving oligodendrocyte cell body and the newly generated, remyelinating myelin sheath. (Frame rate: 2 frames per second).
