## Supplementary figures and images for "Motor Learning Promotes Remyelination via New and Surviving Oligodendrocytes"

### Supplementary Video 3

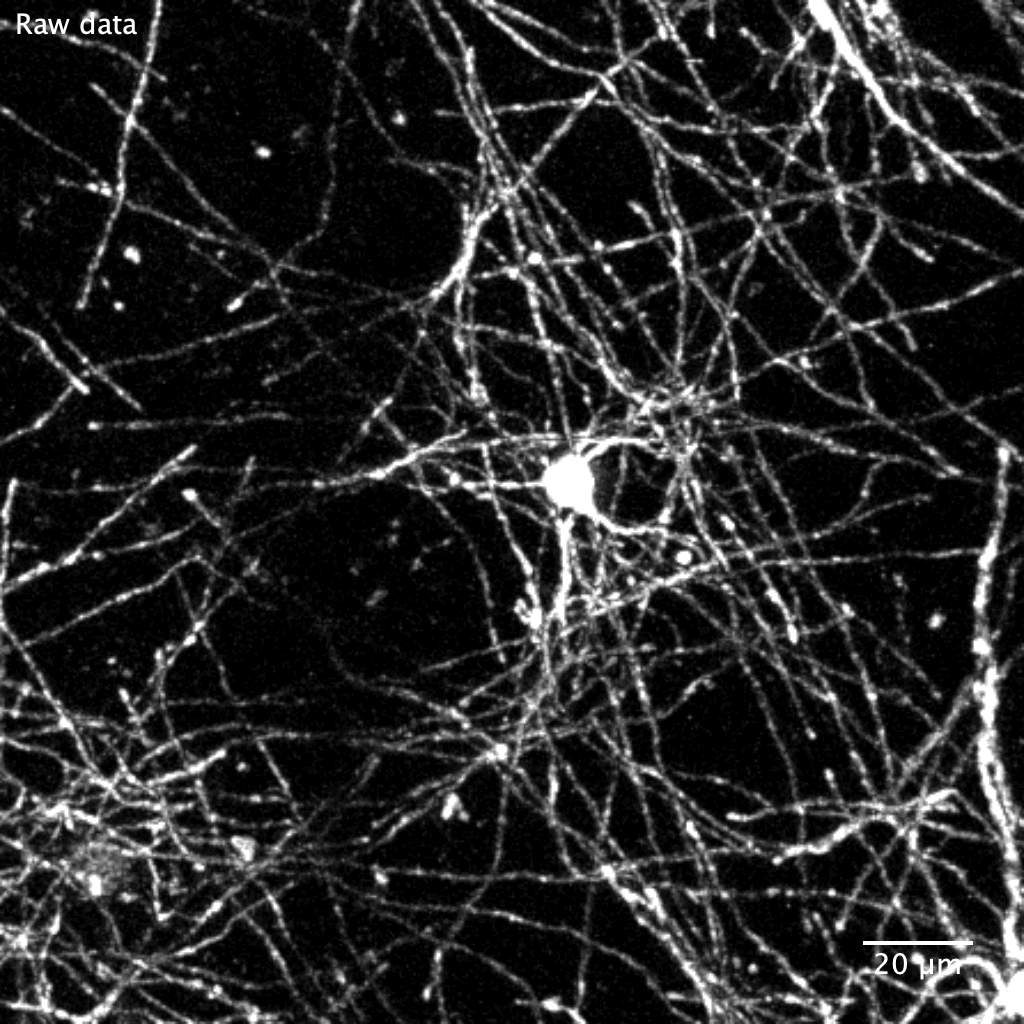

### Supplementary Video 14

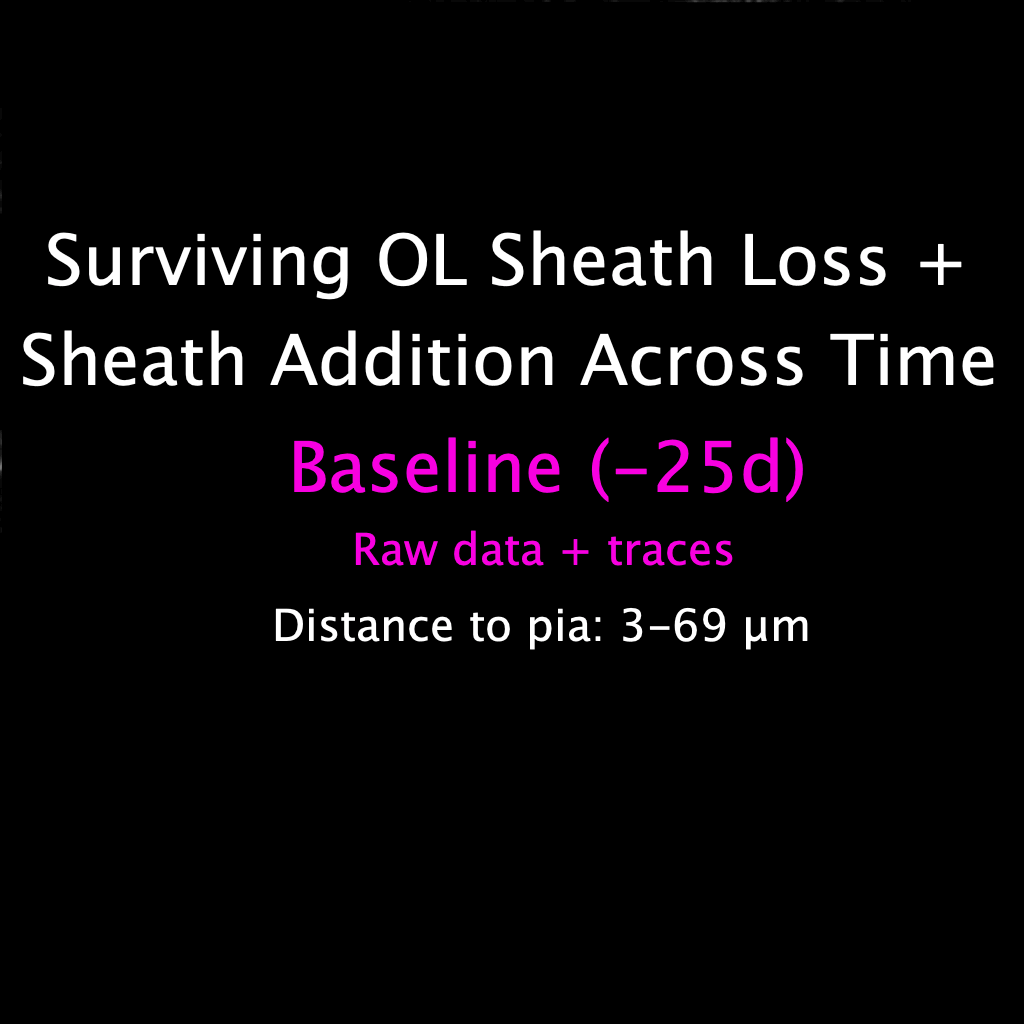
